# Correlation-aware discovery of co-occurring mutational signatures in cancer

**DOI:** 10.64898/2026.09.14.751548

**Authors:** Hu Jin, Benedikt Geiger, Dominik Glodzik, Doga C Gulhan, Peter J Park

**Author notes:** These authors contributed equally: Hu Jin, Benedikt Geiger.

## Abstract

Somatic mutations in cancer genomes record the activities of diverse mutational processes. Mutational signature analysis has advanced mechanistic understanding of mutagenesis and informed clinical decision-making, yet existing methods assume independence among signatures—an unrealistic assumption that can produce composite or contaminated signatures, reduce detection power, and yield inconsistent results. Here we present Cornet (CORrelated NMF ExTraction), a framework for mutational signature discovery that explicitly models co-occurring processes and jointly infers signatures and their correlation structure. Benchmarking on simulated data shows Cornet more accurately recovers distinct signatures under strong correlations. Applied to cancer genomes, Cornet enables unsupervised discovery of the colibactin-associated signature SBS88 in oral cancers and identifies the tobacco smoking signature SBS4 in bladder cancer, where it was previously thought absent. Cornet also uncovers a novel mutational process implicated in early-onset colorectal cancer and a signature arising from the interplay between tobacco smoking and ERCC2-mutation-driven nucleotide-excision repair deficiency. Together, these results demonstrate that modeling correlations among mutational processes is essential for high-resolution signature discovery and dissecting the mutational etiology of human cancer.

## Main

Cells in the human body continuously accumulate somatic mutations. Some of these mutations confer a growth advantage, leading to clonal expansion and, ultimately, malignant transformation. The landscape of somatic mutations in cancer is shaped by the combined effect of endogenous (e.g., defective DNA repair) and exogenous (e.g., tobacco smoking) processes acting within the same tumor. Elucidating these mutational processes is a central goal of cancer research [1]. Mutational signature analysis has emerged as a key approach for achieving this goal [2–4]. By applying pattern-recognition algorithms to large cancer genome sequencing datasets, it identifies characteristic mutation patterns and their sequence contexts—mutational signatures—that reflect the underlying mutational processes [5, 6]. These signatures have become a powerful resource for uncovering the mechanisms of mutagenesis and guiding clinical decision-making [2–4, 7].

*De novo* mutational signature discovery is challenging because each tumor represents a cumulative imprint of multiple mutational processes. It thus relies on variation in the activities of these processes across large, heterogeneous tumor cohorts. Most existing computational methods for this task employ non-negative matrix factorization (NMF) or its extensions [8–12]. These have enabled systematic discoveries of mutational signatures and the construction of curated reference catalogs such as COSMIC and Signal [10, 13–15]. However, these methods fundamentally assume independence among signatures. They decompose the mutation count matrix into the product of two lower-rank matrices corresponding to signatures and exposures, respectively. Probabilistically, this formulation can be interpreted within the broader framework of Poisson factorization models widely studied in the machine learning literature, in which each latent factor contributes independently and additively to the observed mutation counts [16].

In cancer genomes, the independence assumption does not hold: mutational processes frequently co-occur and exhibit strong correlations. Such correlations can arise when multiple processes are associated with the same or intrinsically linked underlying causes (e.g., aging, co-administered drugs, or concurrent alcohol and tobacco consumption). Correlations also arise when a single mutagenic source gives rise to multiple coupled signatures because of the distinct DNA lesions and repair mechanisms involved, as observed for mismatch repair deficiency (MMRD), ultraviolet (UV) radiation, and AID or APOBEC activities.

These correlations pose a major challenge for signature discovery because they effectively reduce sample heterogeneity. As a result, existing methods can yield composite signatures that mix signals from multiple processes, thereby masking their underlying components. A canonical example is APOBEC-mediated mutagenesis, which produces either C*>*T or C*>*G mutations depending on whether the resulting uracil is excised to form an abasic site before replication. The two corresponding signatures, SBS2 and SBS13, were heavily conflated in early COSMIC releases (v1 and v2) owing to their strong correlation and limited sample sizes. Their separation was later achieved only when the whole-genome sequencing (WGS) datasets got sufficiently large (COSMIC v3 onward) and was experimentally validated through uracil glycosylase knockout [17]. Similar refinements and separations into multiple components have occurred for other signatures, including SBS7 (UV), SBS10 (polymerase defects), SBS17 (unknown etiology), SBS22 (aristolochic acid exposure), and SBS40 (unknown etiology). Moreover, a number of signatures in the reference catalog were “cleaned” by removing contamination from confounding signatures as more data became available. Nevertheless, limited cohort sizes and the inability of NMF-based methods to resolve correlated processes remain major bottlenecks [18], suggesting that current reference catalogs may still contain composite or contaminated signatures.

Such composite or contaminated signature discoveries have profound consequences. First, weak signatures may be obscured by contamination from more prevalent ones, hindering the detection of mutational processes associated with sporadic mutagen exposures or rare genetic deficiencies, especially in tumor types with limited data. Second, composite or contaminated signatures may appear inconsistent across datasets despite reflecting the same underlying processes, because dataset-specific correlation structures impede *de novo* discovery methods that assume independence. If uncorrected, such algorithmmic variation can confound genuine biological differences, such as tissue-specific signatures [10]. Finally, composite signatures can obscure mechanistic interpretation. For example, SBS40 was initially thought to be ubiquitous across many tumor types, but a recent analysis of a large, geographically diverse kidney cancer cohort resolved SBS40 into three distinct components, revealing that only one is ubiquitous while the others are largely kidney-specific [19].

Here, we introduce Cornet, a correlation-aware framework for mutational signature discovery. Unlike existing NMF-based methods, Cornet removes the independence assumption. Instead, it jointly models and infers signatures and their correlation structure from mutation count data. Through simulation studies and real-data analyses, we demonstrate that Cornet achieves higher sensitivity in discovering non-composite, uncontaminated signatures, even under strong correlations and limited sample sizes. Applying Cornet across multiple tumor types, we uncover several biological insights, including the first unsupervised discovery of the colibactin-associated signature SBS88 in oral cancers and identification of the canonical tobacco smoking signature SBS4 in bladder cancer, where it was previously thought absent. Cornet also resolves previously composite signatures into distinct signatures, uncovering a novel mutational process implicated in early-onset colorectal cancer and another process arising from the interplay between tobacco smoking and ERCC2 mutations, which impair nucleotide-excision repair.

## Results

### Correlated mutational processes hinder accurate signature discovery

Correlations among mutational processes pose a fundamental challenge for accurate signature discovery (**Fig. 1a**). To illustrate this problem, we simulated a dataset of 100 samples influenced by three mutational processes (**Fig. 1b**). Two of these processes, SBS2 and SBS13, correspond to APOBEC mutagenesis and had correlated exposures, i.e., contributing to similar mutational burdens. The third process, SBS5, was independent of SBS2 and SBS13. When applied to the simulated data, NMF recovered three signatures resembling the ground truth, but with substantial cross-contamination. In particular, NMF was unable to cleanly separate the two correlated APOBEC signatures, yielding an SBS13 solution strongly contaminated by SBS2 (**Fig. 1b**). This problem in signature discovery also leads to underestimated correlations of the NMF solutions of SBS2 and SBS13, because the SBS2-like contaminating component in the SBS13 solution “steals” exposure attributions away from the SBS2 solution (**Fig. 1b**). Furthermore, although SBS5 was simulated to be independent of SBS2 and SBS13, its NMF solution also suffered from a similar contamination problem (**Fig. 1b**). Thus, correlations among mutational processes not only distort correlated signatures, but can also propagate contamination to otherwise unrelated ones.

**Figure 1.**
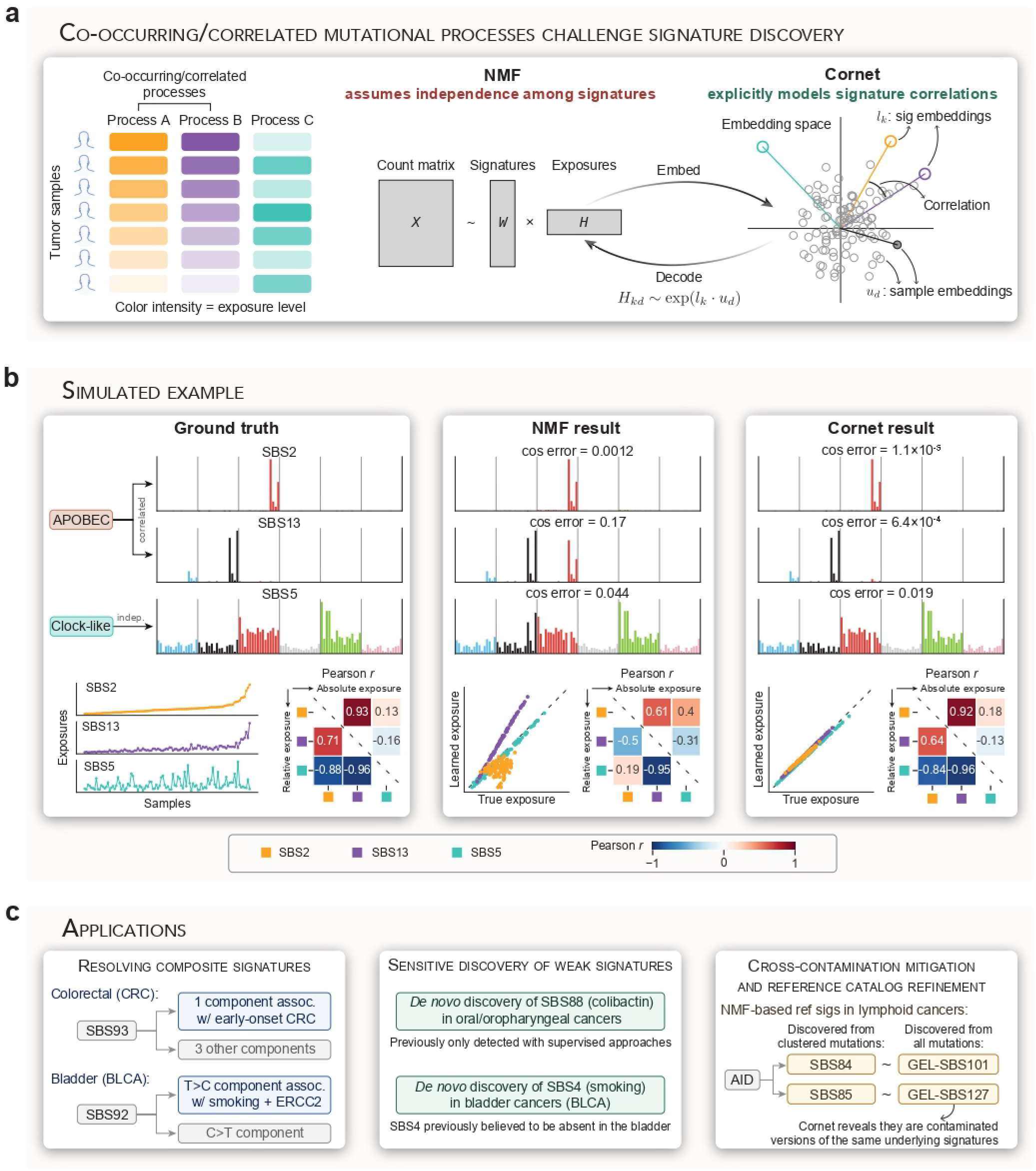
Cornet: a correlation-aware framework for mutational signature discovery. **(a)** Correlated mutational processes are pervasive in cancer—multiple processes frequently co-occur across tumor samples (left), violating the independence assumption underlying existing NMF-based signature discovery methods (center). Cornet addresses this challenge by explicitly modeling correlations in a latent embedding space and jointly inferring signatures and their correlation structure (right). **(b)** Simulated example illustrating the consequences of violating independence. When correlated mutational processes are present (APOBEC-induced SBS2 and SBS13), NMF produces cross-contaminated signatures and erroneous correlation estimates. Cornet accurately recovers all three signatures and their correlation structure. Exposure line plots show ground-truth activities across samples sorted by SBS2 exposure; correlation coefficients are computed from either absolute or L1-normalized relative exposures. **(c)** Summary of Cornet application scenarios and key discoveries presented in this paper.

This simulated data serves only as a simplified illustration of a broader, more complex problem that we aim to solve (**Fig. 1a, c**). In simulations, these cross-contaminations are readily identifiable because the ground truth is known, and the confounded solutions can be corrected by *post hoc* matching to a reference catalog. In real datasets, however, ground truth is unknown, and it is thus difficult to determine whether *de novo* signatures are contaminated or not. As a result, composite signatures containing multiple sub-components may be obtained, and weak novel signatures may be masked by contaminations from more dominant signatures. In the following sections, we demonstrate that these phenomena are widespread in cancer genomes and can hinder downstream analysis and interpretation (**Fig. 1c**).

### Correlation-aware mutational signature discovery with Cornet

We introduce Cornet, a correlation-aware framework for mutational signature discovery based on a revised version of the correlated NMF model [20] (**Methods, Supplementary Methods**). Cornet drops the independence assumption among discovered signatures and instead explicitly models their correlation structure. This is achieved by representing signatures and samples as latent vectors in a shared embedding space, where angles between signature embeddings model correlations among mutational processes, and angles between signature and sample embeddings model the exposures (**Fig. 1a**). These latent representations, together with the signature matrix, are jointly inferred from the mutation count matrix by maximizing the posterior distribution.

When applied to the same simulated data, Cornet accurately recovered the three underlying signatures without the cross-contamination observed in NMF solutions, yielding higher cosine similarities to the ground truth (**Fig. 1b**). Moreover, this clean signature deconvolution was accompanied by accurate inference of the correlation structure among the discovered signatures (**Fig. 1b**).

To benchmark Cornet systematically, we simulated generic datasets with three random signatures (**Methods, Supplementary Fig. S1**). A correlation r (-1≤r≤1) between two signatures (Sig1 and Sig2) was introduced, and the third signature (Sig3) was simulated to be independent. NMF and Cornet were then applied to recover signatures from the simulated mutation count matrices.

When the correlation was negative or weakly positive (r≤0.5), NMF recovered all three signatures with reasonable accuracy (median cosine error*<*0.006). However, when Sig1 and Sig2 were strongly positively correlated, NMF produced increasingly inaccurate solutions (median cosine error 0.032-0.13 for 0.8*<*r≤0.98) (**Supplementary Fig. S1a**). Although these cosine errors appear modest, they correspond to substantial cross-contaminations: a median of 20-41% of weights were attributed to cross-contaminations when NMF solutions were decomposed into ground-truth signatures using non-negative least squares (NNLS) (**Supplementary Fig. S2**). By contrast, Cornet accurately recovered all three signatures across the same correlation range (median cosine error 0.0039-0.033 for 0.8*<*r≤0.98) (**Supplementary Fig. S1a**), with markedly reduced cross-contaminations (2-11%) (**Supplementary Fig. S2**). Cornet produced large errors—while still achieving higher accuracy than NMF—only when correlations were so strong (r*>*0.98) that Sig1 and Sig2 effectively behaved as a single signature.

Cornet also accurately inferred the correlation structure of discovered signatures, whether based on absolute or L1-normalized exposures, except at extreme correlations (**Supplementary Fig. S1b-c**). By comparison, NMF systematically underestimated correlations between Sig1 and Sig2 and frequently inferred incorrect correlation signs. NMF-based correlation estimates involving the independent signature (Sig3) were similarly confounded.

Together, these results demonstrate that Cornet substantially improves both signature discovery and correlation inference from synthetic data, even in the presence of strong correlations among mutational processes.

### Cornet enables sensitive discovery of the colibactin signature SBS88 from small colorectal and oral/oropharyngeal cancer datasets

As a proof of principle using real data, we demonstrate that Cornet can sensitively discover the weak signature SBS88 from smaller datasets than previously required. SBS88 is an experimentally validated signature attributed to colibactin, a genotoxin produced by *pks*+ E. *coli* present in the human large intestine (**Fig. 2a**) [21, 22]. Recent studies using NMF-based algorithms discovered SBS88 in colorectal cancer (CRC) only from large WGS cohorts (n=496, 981, and 2,023 in [22–24], respectively), suggesting that colibactin exposure contributes to CRC tumorigenesis (**Fig. 2b**). Earlier efforts using smaller datasets, however, failed to discover this signature.

**Figure 2.**
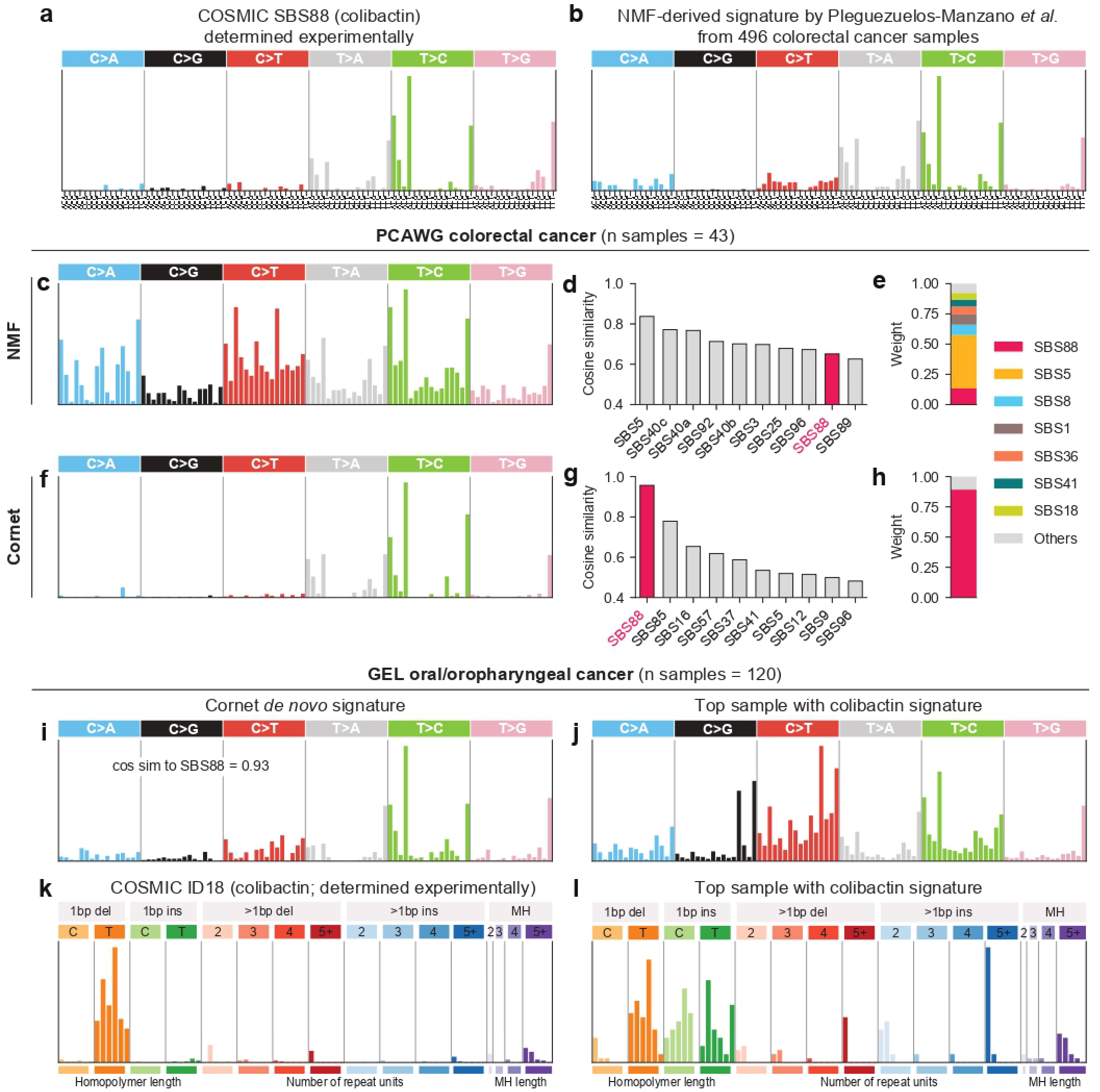
Cornet enables sensitive discovery of the colibactin signature SBS88 from small colorectal and oral/oropharynxgeal cancer datasets. **(a)** COSMIC SBS88, originally determined experimentally. **(b)** NMF signature derived in [22] from 496 colorectal cancer (CRC) samples, resembling SBS88 in (a). **(c)** NMF-derived *de novo* signature from the PCAWG CRC dataset (n=43) with the highest cosine similarity to SBS88. **(d)** Top ten matched COSMIC signatures together with the corresponding cosine similarities for the NMF-derived signature in (c). **(e)** NNLS weights of decomposing the NMF-derived signature in (c) into CRC-relevant reference signatures. **(f-h)** Same as (c-e) but for the Cornet-derived *de novo* signature from the same dataset. **(i)** Cornet-derived *de novo* signature from GEL oral/oropharyngeal cancers (n=120) resembling SBS88. **(j)** SBS mutational spectrum of the GEL oral/oropharyngeal sample with the largest exposure to the *de novo* signature in (i). **(k)** Colibactin-associated COSMIC indel signature ID18, originally determined experimentally. **(l)** Indel mutational spectrum of the same sample in (j).

One such effort was by the PCAWG consortium [14, 25]. The PCAWG cohort included only 43 CRC samples after excluding those with mismatch-repair deficiencies. We repeated *de novo* signature discovery from these 43 samples using both NMF and Cornet. Indeed, NMF failed to discover SBS88. The closest NMF solution was heavily contaminated, despite showing characteristic peaks of T*>*N substitutions at ATA, ATT, and TTT (**Fig. 2c**). When compared to the COSMIC catalog, it matched SBS5 most closely (cosine similarity = 0.84), whereas SBS88 ranked 9^th^ (cosine similarity = 0.65, **Fig. 2d**). Decomposition into CRC-relevant signatures with NNLS showed only 13% contribution from SBS88, with the rest of the signal coming from other more prevalent signatures such as SBS1, SBS5, SBS8, and SBS36 (**Fig. 2e**). Even forcing NMF to output a larger number of signatures did not recover SBS88 (**Supplementary Fig. S3**).

By contrast, Cornet accurately discovered SBS88 from the same 43 CRC samples (cosine similarity = 0.96, **Fig. 2f**). Notably, the Cornet solution was the most similar to SBS88 among all COSMIC signatures (**Fig. 2g**) and barely contaminated, with 89% of contribution coming from SBS88 (**Fig. 2h**).

To test the sensitive discovery of SBS88 more broadly, we applied Cornet to oral/oropharyngeal cancers from the Genomics England (GEL) cohort (n = 120) [26, 27]. Interestingly, Cornet identified a signature highly similar to SBS88 (cosine similarity = 0.93) (**Fig. 2i**). The sample with the largest exposure of this *de novo* signature showed characteristic T*>*N peaks at ATA, ATT, and TTT in its SBS spectrum (**Fig. 2j**), and an indel spectrum with single T deletions at T homopolymers, consistent with the experimentally determined colibactin indel signature ID18 (**Fig. 2k-l**). These results thus indicate that SBS88 is present in oral/oropharyngeal cancers and suggest a potential contribution of colibactin-producing bacteria to tumorigenesis in this tissue.

To the best of our knowledge, this is the first unsupervised *de novo* discovery of SBS88 in oral/oropharyngeal cancers. Previous studies either missed this signature with NMF-based discoveries from the same cohort [15] or relied on the experimentally defined SBS88 for informed detections from independent cohorts via direct comparison, refitting, or dedicated machine learning classifiers [22, 28, 29].

Together, these results demonstrate the potential of Cornet to disentangle weak signatures from more prevalent ones, even in small datasets and without prior knowledge.

### Cornet discovers the canonical AID signatures SBS84 and SBS85 in lymphoid tumors without requiring an analysis restricted to clustered mutations

We next analyzed B-cell non-Hodgkin lymphomas (BNHL) from PCAWG (n = 107). In these malignancies arising from post-germinal center B cells, activation-induced cytidine deaminase (AID) contributes to mutagenesis. Three known signatures are associated with AID: SBS9, corresponding to non-canonical AID activity, leads to widespread mutations along the entire genome and is relatively easy to discover [13, 14]; SBS84 and SBS85, corresponding to canonical AID activity, produce clustered mutations in confined genomic regions and are more challenging to detect [14, 30] (**Fig. 3a-b**). Accordingly, previous *de novo* discovery of SBS84 and SBS85 relied on analyses restricted to clustered mutations to avoid being overwhelmed by non-clustered mutational signatures [14, 30].

**Figure 3.**
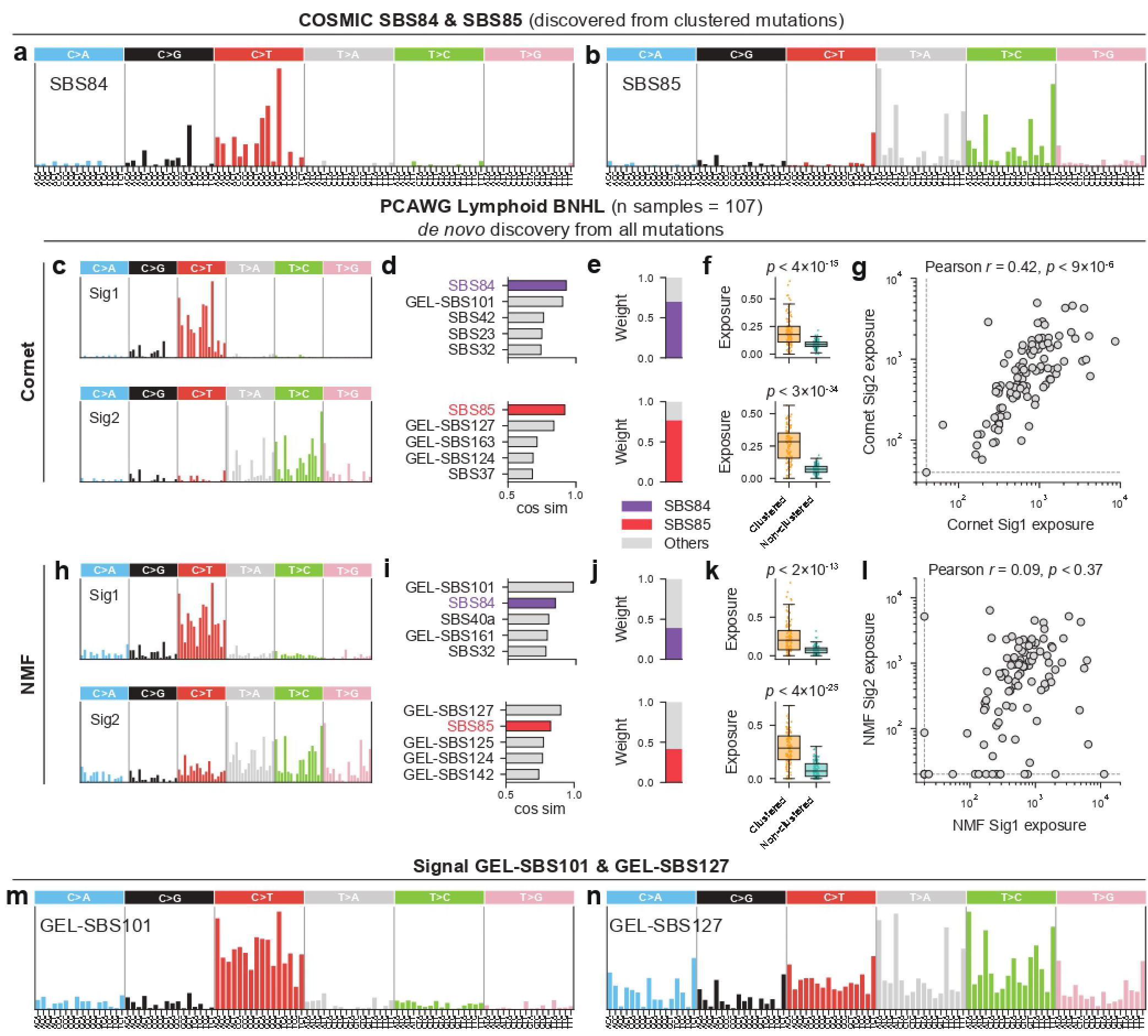
Cornet discovers SBS84 and SBS85 without requiring an analysis restricted to clustered mutations and identifies two likely redundant reference signatures arising from NMF artifacts. **(a-b)** COSMIC SBS84 and SBS85, originally discovered by analyzing clustered mutations in lymphoid tumors with NMF. **(c)** Two *de novo* signatures resembling SBS84 and SBS85, discovered by analyzing all mutations, regardless of whether they are clustered or non-clustered, in PCAWG lymphoid BNHL samples with Cornet. **(d)** Top five matched reference signatures and the corresponding cosine similarities for the two *de novo* signatures in (c). **(e)** NNLS weights of decomposing the two *de novo* signatures in (c) into reference signatures known to be present in BNHL. **(f)** Exposures of the two *de novo* signatures in (c) in clustered vs. non-clustered mutations. **(g)** Exposures of the two *de novo* signatures in (c) plotted against each other. Pearson correlation coefficients and p-values are annotated. For visualization, data points smaller than the dashed lines were plotted on the dashed lines, without affecting the correlation calculations. **(h-l)** Same as (c-g) but for the two NMF-derived *de novo* signatures. NMF-based discovery was also performed using all mutations, regardless of whether they are clustered or non-clustered. The two *de novo* signatures with the highest similarities to COSMIC SBS84 and SBS85, respectively, are shown. **(m)** GEL-SBS101, a reference signature in the Signal catalog, originally discovered by NMF exclusively from GEL lymphoid tumors. **(n)** GEL-SBS127, a reference signature in the Signal catalog, originally discovered by NMF from GEL lymphoid tumors as well as several other tumor types.

Remarkably, Cornet discovered two signatures resembling SBS84 and SBS85 (cosine similarity = 0.93 and 0.92) while using all mutations as input, irrespective of whether they were clustered or nonclustered (**Fig. 3c**). When compared to reference signatures from COSMIC (denoted as SBSn) and Signal (mostly GEL-derived, thus denoted as GEL-SBSn), these two *de novo* signatures were most similar to SBS84 and SBS85 (**Fig. 3d**). Decomposition into BNHL-relevant signatures confirmed they predominantly captured SBS84 and SBS85, with only minor contaminations (**Fig. 3e**). Consistent with their biological interpretation, both signatures were significantly enriched in clustered mutations and correlated with each other as expected (**Fig. 3f-g**).

By contrast, NMF failed to robustly recover SBS84 and SBS85 without restricting the analysis to clustered mutations. The two NMF solutions closest to SBS84 and SBS85 showed lower cosine similarities (0.85 and 0.83), severe contamination by other signatures, weaker enrichment in clustered mutations, and lacked the expected correlation (**Fig. 3h-l**). Instead, these solutions were most similar to GEL-SBS101 and GEL-SBS127 (cosine similarities = 0.98 and 0.90), reference signatures originally discovered by applying NMF to all mutations irrespective of clustering in lymphoid tumors (GEL-SBS101) or lymphoid and other tumor types (GEL-SBS127) (**Fig. 3m-n**) [15]. Although considered novel in [15] due to their lack of similarity to known reference signatures, a side-by-side comparison of the NMF and Cornet results suggests that these signatures represent SBS84 and SBS85 entangled with other contaminating signals (**Fig. 3h-l**, compared with **Fig. 3c-g**). Together, these findings demonstrate that failure to resolve correlated subcomponents can cause composite NMF signatures to be misinterpreted as novel, and that Cornet can help identify and resolve such mixtures.

### Cornet resolves SBS93 into four constituent component signatures in colorectal cancers

We next applied Cornet to a larger CRC cohort comprising 1,592 mismatch-repair-proficient (MMRP) samples from GEL [26, 27]. In total, fifteen *de novo* signatures were discovered, nine of which matched CRC-associated reference signatures, including SBS1 (aging), SBS2/13 (APOBEC), SBS17a/b (ROS damage), SBS18 (ROS damage), and SBS88 (colibactin). (**Supplementary Fig. S4a**). We therefore focused on the remaining six signatures. To distinguish Cornet-derived signatures from reference signatures, we prefixed them with “CRC-”. Four of these six matched SBS89, SBS93, GEL-SBS121, and GEL-SBS128, and were named CRC-SBS89, CRC-SBS93, CRC-SBS121, and CRC-SBS128, respectively, while the remaining two were designated CRC-SBS-A and CRC-SBS-B (**Supplementary Fig. S4a**).

SBS93 is a signature of unknown etiology originally discovered by NMF in gastric and esophageal cancers and later reported in CRC, suggesting a mutational process shared in the gastrointestinal tract (**Fig. 4a**) [9, 23, 24]. Cornet confirmed the presence of SBS93 in CRC through unsupervised discovery. Although CRC-SBS93 retained the defining T[T*>*C]A, T[T*>*G]A, and T[T*>*G]T peaks of SBS93, it was markedly sparser with a reduced flat background contribution, consistent with decreased contamination (**Fig. 4b**). Analysis of the correlation structure of Cornet-derived signatures revealed that CRC-SBS93, CRC-SBS-A, CRC-SBS121, and CRC-SBS128 formed a tightly correlated cluster (**Fig. 4b, Supplementary Fig. S4b**). All four signatures were also tightly correlated with the indel signature ID14, previously linked to SBS93 [23], with CRC-SBS93 showing the strongest correlation (**Supplementary Fig. S4c**).

**Figure 4.**
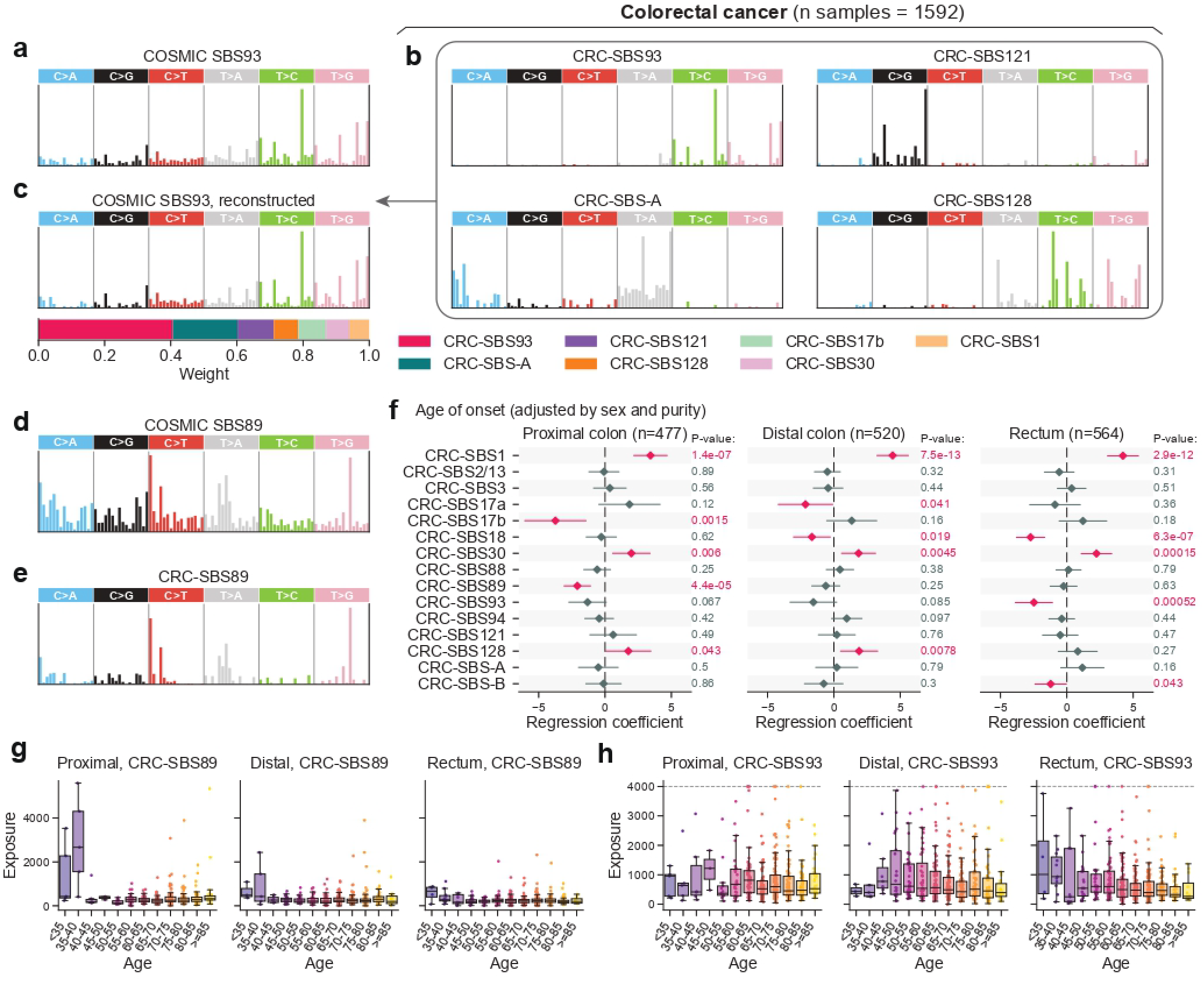
Applying Cornet to colorectal cancers (CRC) resolves SBS93 into four components, refines SBS89, and provides new insights into signatures associated with early-onset CRC. **(a)** COSMIC SBS93, originally discovered by NMF in [9]. **(b)** CRC-SBS93, CRC-SBS-A, CRC-SBS121, and CRC-SBS128, four *de novo* signatures discovered by Cornet that underlie COSMIC SBS93. **(c)** Top: reconstructed COSMIC SBS93 with Cornet-derived *de novo* signatures (cosine similarity to the original COSMIC SBS93 = 0.98). Bottom: normalized weights used in the reconstruction. **(d)** COSMIC SBS89. **(e)** CRC-SBS89 discovered by Cornet. **(f)** Association between age of onset and Cornet-derived mutational signatures. Linear regression of age of onset was performed on the exposures of Cornet-derived *de novo* signatures, with patient sex and tumor purity included as covariates. Proximal colon (left), distal colon (middle), and rectal (right) samples were analyzed separately. **(g)** Exposure of Cornet-derived CRC-SBS89 in different sites and age bins. **(h)** Exposure of Cornet-derived CRC-SBS93 across different anatomical sites and age bins. Exposures *>*4000 are truncated to 4000 (dashed line) to facilitate visualization, but the box plots are unaffected.

These results indicate that COSMIC SBS93 represents a composite of four correlated constituent signatures. Their strong correlations prevented separation by NMF, whereas Cornet successfully unmixed them. Indeed, SBS93 could be accurately reconstructed by these four components, along with minor contributions from additional contaminating signatures (**Fig. 4c**). This unmixing was independently supported by principal component analysis (PCA), in which trinucleotide mutation types associated with each constituent signature formed distinct clusters in PCA space (**Supplementary Fig. S4d**).

Similarly, CRC-SBS89 was sparser than COSMIC SBS89 while preserving its characteristic G[T*>*G]G, A[C*>*T]A, A[C*>*T]T, and C[T*>*A]N peaks, again consistent with reduced contamination (**Fig. 4d-e**). SBS89 has been reported in both normal and cancerous colon tissues, but its etiology remains unclear [23, 24, 31]. Interestingly, CRC-SBS89 was tightly correlated with the indel signature ID23 known to be caused by aristolochic acid exposures (**Supplementary Fig. S4c**), and its C[T*>*A]N peaks resembled those of the aristolochic acid signature SBS22a. It is thus likely that similar exogenous exposures underlie CRCSBS89, although future studies are needed to validate this connection. Finally, CRC-SBS-B is a novel signature showing similarity to a recently reported signature in CRC [32] (**Supplementary Fig. S4a**).

### Unmixing of SBS93 reveals its association with early-onset colorectal cancers

The incidence of early-onset CRCs, defined as those diagnosed before age 50, has increased globally over the past two decades, yet its underlying causes remain poorly understood [33, 34]. To identify mutational processes associated with age of onset, we regressed patient age at diagnosis against exposures of Cornet-derived signatures in the GEL CRC cohort, controlling for sex and tumor purity (**Fig. 4f**). Site-specific analyses were performed for proximal colon, distal colon, and rectum, separately.

Among all anatomical sites, CRC-SBS1 and CRC-SBS30 increased with later age at diagnosis, consistent with mutagenic processes that accumulate over time. By contrast, several signatures were associated with early onset in a site-specific manner, including those linked to reactive oxygen species (ROS) damage: CRC-SBS17b was associated with early onset in proximal colon cancers, whereas CRC-SBS18 was associated with early onset in rectal cancers, suggesting distinct local sources of ROS contributing to early tumorigenesis. Notably, SBS17 has recently been linked to ethanol and acetaldehyde in rodent models [35], raising the possibility that alcohol consumption contributes to early-onset CRC.

CRC-SBS89 showed a strong association with early onset in proximal colon cancers. Patients diagnosed before age 40 were particularly enriched for this signature, with an average of more than 2,000 mutations attributable to CRC-SBS89, compared to approximately 300 mutations in patients diagnosed after age 40 (**Fig. 4g**). A similar trend was observed in distal colon cancers, although limited sample size precluded statistical significance in the regression analysis (**Fig. 4f-g**). Together with evidence implycating exogenous mutagens such as aristolochic acid (see previous section), these findings suggest that environmental exposures, potentially via diet or folk medicine, may contribute to very early-onset proximal colon cancers.

Among the four constituent components of SBS93 identified by Cornet, only CRC-SBS93 itself was associated with early onset, specifically in rectal cancers. Unlike CRC-SBS89, CRC-SBS93 showed a gradual decrease with age of onset (**Fig. 4h**). In distal colon cancers, CRC-SBS93 increased with younger age down to age 40, but patients diagnosed before age 40 had lower exposures, resulting in a non-linear relationship and a non-significant association in the linear regression analysis (**Fig. 4f, h**). Excluding patients younger than 40 restored a significant association (**Supplementary Fig. S5**). In proximal colon cancers, CRC-SBS93 showed minimal age-related variation (**Fig. 4h**).

Together, these results identify three classes of mutational processes potentially contributing to early-onset CRC: site-specific ROS-associated damages (CRC-SBS17b and CRC-SBS18), CRC-SBS89 in proximal colon cancers, and CRC-SBS93 in distal colon and rectal cancers. While a prior study reported age-dependent decreases of SBS89 and SBS93 using composite COSMIC signatures [23], Cornet resolves these effects to their precise underlying components, revealing site-specificity and, in some cases, opposite associations. Notably, only one of the four SBS93 components was linked to early onset, whereas another component CRC-SBS128 was associated with later onset (**Fig. 4f**), underscoring the limitations of composite signatures. Finally, although SBS88 has been reported to be enriched in earlyonset CRC in an independent and geographically diverse cohort [24], we did not observe this association in the GEL cohort, possibly reflecting limited geographic variation in colibactin exposure within this single-country cohort.

### Cornet resolves tobacco smoking-associated signatures in bladder cancers

Tobacco smoking is a well-established risk factor for bladder cancer (BLCA) [36]. However, the canonical tobacco smoking-associated signature SBS4—characterized by C*>*A and T*>*A substitutions—has not been detected in BLCA despite its prevalence in other smoking-related tumor types such as lung and liver [14, 37]. Instead, SBS92, a distinct signature characterized by T*>*C and C*>*T substitutions, was recently discovered in BLCA using NMF and found to be enriched in smokers (**Fig. 5a**) [9], suggesting tissue-specific tobacco smoking-associated mutagenesis in the bladder.

**Figure 5.**
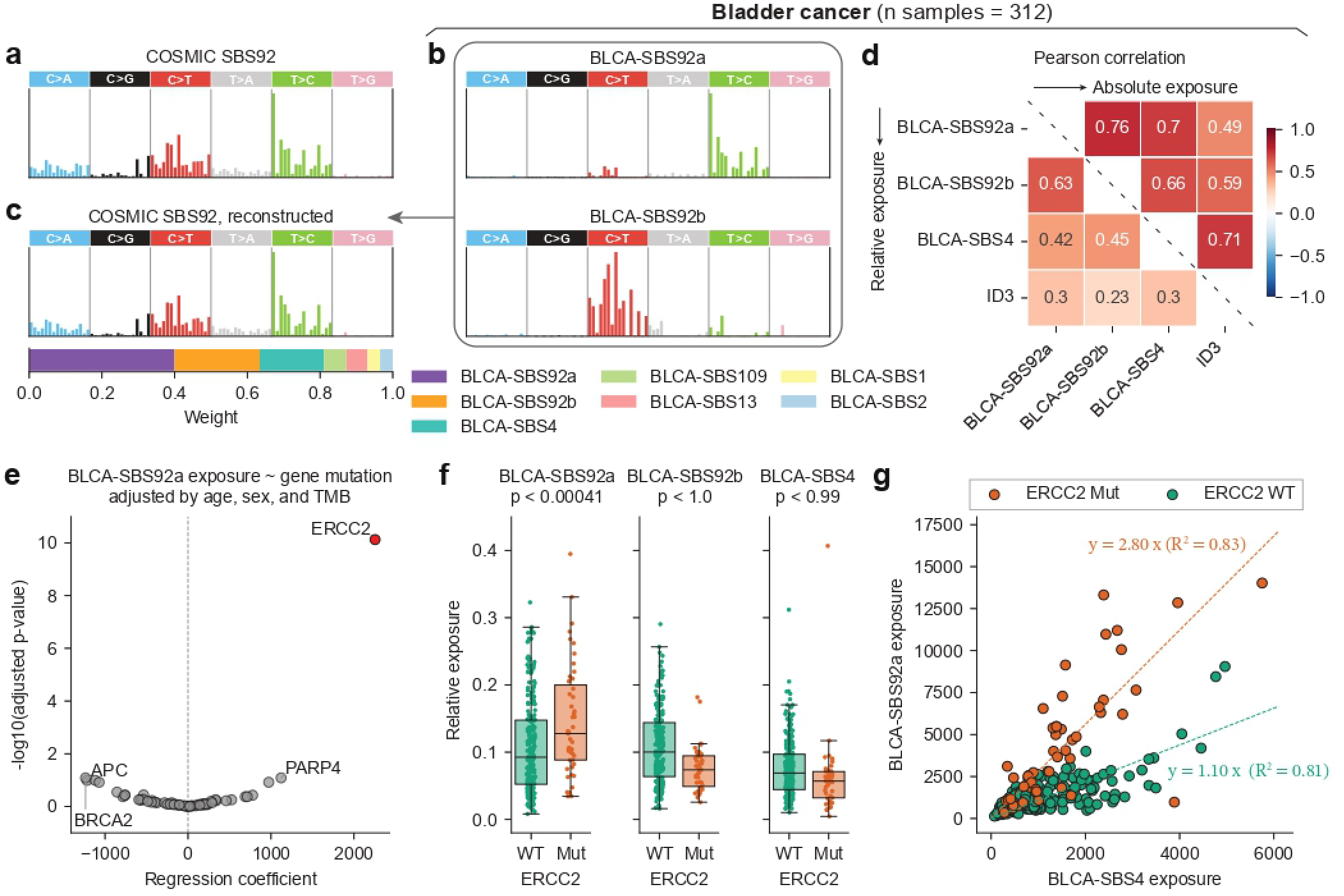
Cornet resolves SBS92 into two components in bladder cancers (BLCA), and the T*>*C component is associated with tobacco smoking combined with ERCC2 mutations. **(a)** COSMIC SBS92, originally discovered by NMF from BLCA in [9]. **(b)** BLCA-SBS92a and BLCA-SBS92b, two *de novo* signatures discovered by Cornet that underlie COSMIC SBS92. **(c)** Top: reconstructed COSMIC SBS92 using Cornet-derived *de novo* signatures (cosine similarity to the original COSMIC SBS92 = 0.99). Bottom: normalized weights used in the reconstruction. **(d)** Pearson correlation coefficients between the exposures of BLCA-SBS92a, BLCA-SBS92b, BLCA-SBS4, and ID3. The upper triangle shows correlations calculated using absolute exposures. The lower triangle shows correlations calculated using L1-normalized relative exposures. **(e)** Volcano plot for the linear regression of BLCA-SBS92a exposures against gene-level mutation status. Patient age, sex, and tumor mutational burden (TMB) were included as covariates in the regression. Genes with an adjusted p-value *<*0.05 are colored red. **(f)** L1-normalized relative exposures of BLCA-SBS92a, BLCA-SBS92b, and BLCA-SBS4, comparing ERCC2-widetype (WT) and ERCC2-mutant (Mut) samples. One-sided p-values from the t-test are annotated. **(g)** Absolute exposures of BLCA-SBS92a vs. BLCA-SBS4 for ERCC2-widetype (WT) and ERCC2-mutant (Mut) samples separately. Linear regression slopes are annotated. One outlier sample with BLCA-SBS4 exposure more than four times higher than that of any other sample was removed.

We applied Cornet to a combined cohort of 312 WGS BLCA samples from PCAWG and GEL [25– 27]. As in prior analyses, we prefixed Cornet-derived signatures with “BLCA-” for clarity. Notably, a signature resembling SBS4 was discovered, sharing the characteristic C[C*>*A]N and C[T*>*A]N peaks, and was therefore denoted BLCA-SBS4 (cosine similarity to SBS4 = 0.84, **Supplementary Fig. S6a**). In addition, Cornet separated SBS92 into two tightly correlated signatures capturing its T*>*C and C*>*T components, respectively (BLCA-SBS92a and BLCA-SBS92b, **Fig. 5b**). Together with other contributing signatures such as BLCA-SBS4, these two components accurately reconstructed the SBS92 spectrum, indicating that the NMF-derived SBS92 is a composite signature (**Fig. 5c**). All three signatures—BLCA SBS4, BLCA-SBS92a, and BLCA-SBS92b—were significantly correlated with each other and with the tobacco smoking-associated indel signature ID3 (**Fig. 5d**).

To gain insights into why multiple tobacco smoking-associated signatures are present in BLCA, we examined associations between signature exposures and gene-level mutations while controlling for patient age, sex, and total mutational burden. Interestingly, BLCA-SBS92a exposure was strongly and specifically elevated in tumors harboring ERCC2 mutations, whereas BLCA-SBS92b and BLCA-SBS4 showed no comparable gene associations (**Fig. 5e-f, Supplementary Fig. S6b**). Stratifying tumors by ERCC2 mutation status revealed a linear relationship between BLCA-SBS92a and BLCA-SBS4 exposures in both subsets, but with different slopes: approximately 1:1 in ERCC2-widtype tumors and 3:1 in ERCC2-mutant tumors (**Fig. 5g**). By comparison, the relationship between BLCA-SBS92b and BLCA-SBS4 was similar in ERCC2-wildtype and -mutant tumors (**Supplementary Fig. S6c**).

Together, these results show that Cornet refines the landscape of tobacco smoking-associated muta tional signatures in BLCA by uncovering the presence of the canonical SBS4 and decomposing SBS92 into two biologically distinct components. In particular, BLCA-SBS92a represents the combined effect of smoking-induced DNA damage and impaired nucleotide excision repair (NER), and is selectively elevated in tumors with ERCC2 mutations.

## Discussion

By explicitly modeling correlations between signatures, Cornet improves the power to resolve distinct mutational processes while mitigating contamination and compositional mixing that can arise in NMF-based analyses. Leveraging this improved sensitivity, we gained several insights that improve our current understanding of mutational signature landscapes in human cancer.

A particular illustrative example is tobacco smoking-associated mutagenesis in bladder cancer (BLCA). Contrary to the prevailing view that the canonical tobacco signature SBS4 is absent in BLCA, Cornet identified SBS4 alongside SBS92. Recent studies have further shown that SBS92, although initially identified in BLCA, is also present in other tobacco smoking-related tumor types, including lung, liver, and head and neck cancers [15, 38, 39]. Together, these findings suggest that differences in tobacco smoking-associated mutagenesis across tumor types are primarily quantitative rather than qualitative: both SBS4 and SBS92 occur broadly, but BLCA is enriched for SBS92 whereas other tumor types are enriched for SBS4. This quantitative shift may reflect organ-specific differences in exposure to tobacco-derived metabolites, with most compounds metabolized in the liver and end products excreted through the bladder, or differences in DNA repair pathways involved in resolving smoking-induced lesions.

Consistent with the latter possibility, we observed that ERCC2 mutations are specifically associated with elevated exposures of BLCA-SBS92a, the T*>*C component of SBS92. ERCC2 is recurrently mutated in BLCA but rarely in other tumor types, implicating ERCC2-dependent NER as a key modulator of tobacco smoking-associated mutagenesis in the bladder. Because NER deficiency resulting from ERCC2 mutations confers sensitivity to cisplatin and related agents, strong BLCA-SBS92a exposure may serve as a clinically informative biomarker [40–42]. Notably, the clock-like signature SBS5 has previously been linked to tobacco smoking, particularly in BLCA [37]. Our results suggest that this association likely arose because the spectrum of SBS5 resembles that of SBS92, which was unknown at the time due to small sample sizes and limited power of NMF-based signature discovery.

As cancer genome sequencing efforts continue to expand [26, 43–45], additional signatures are being discovered and added to reference catalogs. However, because most reference signatures (apart from those identified directly from experiments) have been extracted using NMF-based methods, they may suffer from NMF-related algorithmic artifacts, including cross-contaminations and inter-mixing. Our finding that GEL-SBS101 and GEL-SBS127 likely represent algorithmic variants of SBS84 and SBS85 exemplifies this issue. Applying Cornet to existing datasets provides a principled strategy to identify such redundancies, clean contaminating signals, and decompose composite signatures that may still remain in reference catalogs. These refinements will be critical for guiding follow-up etiology studies and for distinguishing methodological artifacts from genuine biological variation across tumor types and disease contexts.

Beyond cancer genomes, mutational signature analysis has become routine in studies of somatic and mosaic mutations in healthy tissues [31, 46–52]. Unlike cancer cohorts, which often exhibit substantial heterogeneity due to high mutational burdens, diverse extrinsic exposures, and various repair deficiencies, datasets derived from healthy tissues are typically more homogeneous, posing a major challenge for conventional NMF-based signature discovery. Cornet may be particularly well-suited to such settings because it is designed to enable robust signal extraction even when variation is limited. We anticipate that Cornet will facilitate future studies of mutational signatures in normal tissues, which have proven valuable for understanding human development and clonal expansion during pre-cancer stages [53–58].

Finally, we view Cornet as complementary to existing methods rather than a replacement. Standard NMF approaches remain valuable for their simplicity and computational efficiency, and our previously developed minimum-volume NMF framework addresses a separate “weight-stealing” problem that is particularly relevant for flat signatures [11]. In contrast, Cornet excels at resolving correlated and composite signatures and mitigating cross-contaminations, especially in homogeneous datasets, smaller cohorts, or specific biological contexts such as healthy tissues. Integrating these complementary approaches, together with other data modalities including epigenetic and clinical information, will be essential for constructing a more accurate and biologically grounded map of mutational processes across tissues and disease states [59, 60].

## Methods

### The Cornet algorithm

Standard non-negative matrix factorization (NMF) provides a widely used framework for mutational signature discovery. However, it treats exposures as deterministic parameters and does not explicitly model correlations between signatures across samples. To overcome this limitation, we introduce Cornet, a correlation-aware framework for mutational signature discovery based on a revised version of the correlated NMF model [20]. Cornet is a probabilistic extension of NMF that models exposures as correlated random variables via a shared latent embedding space.

We first review standard NMF and its probabilistic interpretation in the context of mutational signature discovery. We then introduce the generative model of Cornet and show that it leads to a regularized KL-divergence objective with an additional low-rank structure on the exposure matrix. Model parameters and latent embeddings are estimated via a maximum a posteriori (MAP) approach. We conclude with implementation details and practical considerations.

### Standard NMF

The input data of mutational signature discovery is a *V* -by-*D* nonnegative matrix *X* of mutation counts, where *V* denotes the number of mutation types (e.g., *V* = 96 for standard SBS signatures with trinucleotide contexts) and *D* denotes the number of samples. NMF aims to factorize *X* as *X* ≈ *WH* under the constraint that both *W* and *H* are nonnegative, where *W* is the *V* -by-*K* signature matrix and *H* is the *K*-by-*D* exposure matrix. Here, *K* denotes the number of signatures.

Effectively, NMF represents the mutation spectrum of each sample (a column of *X*) as a linear combination of signatures (columns of *W*), with coefficients given by the corresponding exposure vector (a column of *H*). Given an appropriate quantification of “distance” between *X* and *WH*, such as the generalized Kullback-Leibler (KL) divergence *D*_KL_, this factorization can be obtained by solving the optimization problem

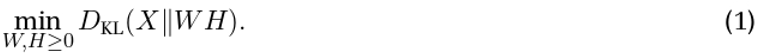

From a probabilistic perspective, the optimization problem in (1) is equivalent to a maximum likelihood approach induced by Poisson noise. Specifically, treating *W* and *H* as model parameters, the corresponding generative model is

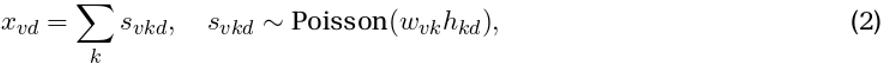

where the latent variable *s*_*vkd*_ represents the number of mutations of type *v* in sample *d* contributed by signature *k*. Assuming conditional independence of *s*_*vkd*_ across all three indices given model parameters, the log-likelihood marginalized over the latent variables *s*_*vkd*_ can be shown to be [61]

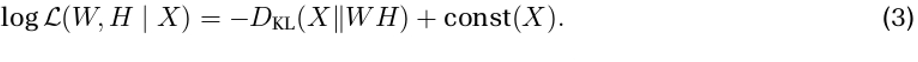

Furthermore, the standard multiplicative update algorithm for solving (1) can be interpreted as an expectation-maximization (EM) algorithm for maximum likelihood estimation under the generative model (2) [61, 62].

We emphasize that in standard NMF, the exposure matrix *H* is treated as a collection of deterministic parameters without a probabilistic model for the exposure vectors *h*_*d*_. As a result, dependencies between signatures are not explicitly modeled, and correlations in their exposures cannot be captured.

### The generative model of Cornet

To explicitly model correlations between signatures, we treat the exposure vectors *h*_*d*_ as random variables. Specifically, we assume that the log-exposures follow a low-dimensional latent Gaussian model

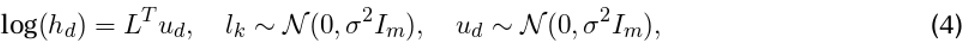

where *L* = (*l*_1_, …, *l*_*K*_) ∈ ℝ^*m×K*^ is a matrix of signature embeddings and *u*_*d*_ ∈ ℝ^*m*^ is a sample embedding. Under this model, log(*h*_*d*_) | *L* ∼ *N* (0, *σ*^2^*L*^*T*^*L*). The covariance structure of the log-exposures is therefore given by

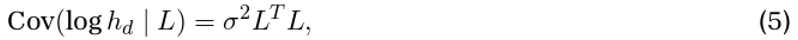

so that correlations between signature exposures are determined by the similarity of their embeddings. In particular, the dot product 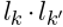 controls the covariance between the exposures of signatures *k* and *k*^*0*^. By symmetry, the model also induces a covariance structure across samples. For a fixed signature *k*, the log-exposure vector across samples satisfies 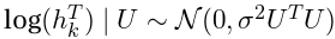, where *U* = (*u*_1_, …, *u*_*D*_) ∈ ℝ^*m×D*^ is the matrix of sample embeddings.

An equivalent generative model is given by

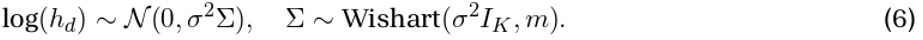

This formulation induces the same marginal distribution of *h*_*d*_ through the factorization Σ = *L*^*T*^*L* [20]. While (6) makes the covariance structure of the log-exposures more explicit, we adopt the embedding formulation in (4) as the primary representation of the generative model because of its more intuitive geometric interpretation. Each signature *k* is associated with an embedding vector *l*_*k*_ ∈ ℝ^*m*^, each sample *d* with a latent vector *u*_*d*_ ∈ ℝ^*m*^, and the exposure of signature *k* in sample *d* is determined by their inner product, log(*h*_*kd*_) = *l*_*k*_ · *u*_*d*_.

Combining this with the Poisson observation model as in standard NMF, the full generative model of Cornet is

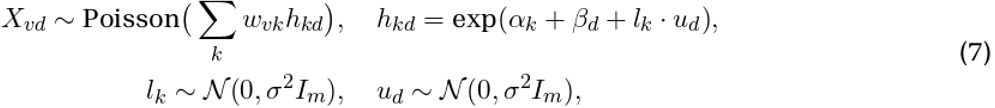

where the additional parameters *α* ∈ ℝ^*K*^ and *β* ∈ ℝ^*D*^ account for global signature and sample effects, respectively. Indeed, the offsets *α* and *β* affect only the mean of the log-exposures, while the covariance structure of the log-exposures is unchanged.

The model parameters are (*W, α, β, σ*^2^), and (*L, U*) are latent variables. During optimization, we enforce that signatures are normalized probability vectors, i.e., 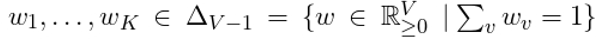.

Although our proposed Cornet model is largely based on the generative process of the correlated NMF model described in [20], there are a few differences:

- We resolve the scaling ambiguity by normalizing the signatures and introducing signature-specific offsets *α*_*k*_. By comparison, in [20], the signatures are not normalized and only sample-specific offsets *β*_*d*_ are introduced.
- We do not introduce priors on the signatures *W*. Instead, *W* is treated as a model parameter and learned directly.
- In our model, the signature and sample embeddings share the same variance parameter *σ*^2^.
- *σ*^2^is not treated as a hyperparameter, as in [20], but is learned during inference.

### Maximum a posteriori estimation

Since the latent variables *L* and *U* make the marginal log-likelihood

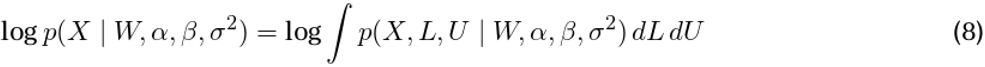

intractable to optimize directly, we optimize the joint log-density

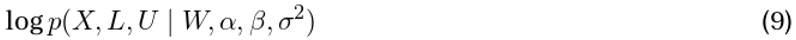

with respect to both the latent embeddings (*L, U*) and the model parameters (*W, α, β, σ*^2^). For fixed model parameters, maximizing this objective with respect to (*L, U*) is equivalent to maximum a posteriori (MAP) estimation of the latent embeddings.

This MAP formulation is closely related to the mean-field variational inference approach of [20], which uses independent point-mass variational distributions for the latent embeddings. In the usual limiting interpretation of such degenerate variational families, this variational treatment yields the same optimization objective as (9). Our overall optimization problem nevertheless differs from that of [20] because we normalize the signatures, omit priors on *W*, introduce signature-specific offsets *α*_*k*_, place the same *σ*^2^-dependent Gaussian prior on both signature and sample embeddings, and learn the variance parameter *σ*^2^.

Under the Cornet generative model, the resulting objective is given by

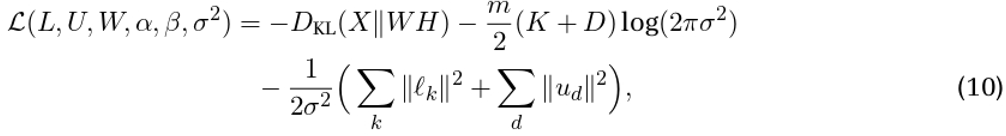

where the first term arises from the Poisson model and the remaining terms arise from the Gaussian priors on the signature and sample embeddings (see **Supplementary Methods**). Thus, Cornet can be viewed as a version of NMF with an additional factorization of the exposure matrix to model correlations, together with a regularization term induced by the Gaussian priors on the latent embeddings.

We optimize (10) by alternating between the following two steps until convergence:

- For fixed model parameters (*W, α, β, σ*^2^), update the latent embeddings (*L, U*) by maximizing (10) with respect to *L* and *U*.
- For fixed latent embeddings (*L, U*), update the model parameters (*W, α, β, σ*^2^) by maximizing (10) with respect to *W, α, β*, and *σ*^2^.

The detailed optimization algorithm is described in **Supplementary Methods**.

### Remarks on software implementation

Cornet is implemented in the Python package Sonata, available at https://github.com/parklab/Sonata. To facilitate the application of Cornet to datasets across a wide range of biological contexts, we implemented Sonata using the popular anndata framework for handling annotated data matrices [63]. For mutational signature discovery, we integrated Cornet within MuSiCal (available at https://github.com/parklab/MuSiCal), a comprehensive framework for mutational signature analysis we previously developed [11]. Specifically, Cornet is integrated within the *de novo* signature discovery pipeline of MuSiCal, which includes parallel execution of signature discovery algorithms across bootstrapped replicates of input data, solution filtering, signature clustering, and automatic selection of the number of signatures *K*.

### Remarks on hyperparameters

Two hyperparameters are involved in the Cornet algorithm: the number of signatures *K* and the embedding dimension *m*. Selection of *K* is handled by the *de novo* signature discovery pipeline of MuSiCal. In short, MuSiCal runs Cornet multiple times with different initializations for each candidate *K*. After filtering out potential bad solutions caused by local optima, MuSiCal selects the best *K* by examining the clustering structure of the solutions from the multiple replicates [11]. The embedding dimension *m* controls the flexibility of the covariance structure, since correlations between signatures are modeled through *L*^*T*^*L*. Smaller values of *m* impose a lower-rank structure and therefore a more restricted covaryance model. In all analyses, we set *m* = *K*, which avoids imposing any additional rank constraint on the covariance structure. Choosing *m > K* is unnecessary, since *K* embedding vectors can span at most a *K*-dimensional space.

### Simulation studies

To compare the performance of Cornet and NMF in disentangling correlated signatures, we constructed synthetic datasets consisting of three SBS signatures, Sig1, Sig2, and Sig3, each randomly drawn from a symmetric Dirichlet distribution. Exposures were simulated according to the generative model in (7) to induce controlled correlation structures between the three signatures. Specifically, the embedding *l*_1_ of Sig1 was fixed to (1, 0, 0)^*T*^, the embedding *l*_3_ of Sig3 was fixed to (0, 0, 1)^*T*^, and the embedding *l*_2_ of Sig2 was taken to be a unit vector in the *x*-*y* plane forming an angle *θ* with *l*_1_, where *θ* ∼ Unif(0, *π*). This ensures that the correlation between the log-exposures of Sig1 and Sig2 varies between −1 and 1, while Sig3 remains independent of Sig1 and Sig2. Mutation count matrices were then sampled from the Poisson distribution as in (7). In total, 1,000 synthetic datasets were simulated independently, each containing 100 samples and around 1,000 mutations per sample on average. Cornet and NMF were then applied to these synthetic datasets. To facilitate comparison, both algorithms were initialized with the same random signatures for a given dataset. Other model parameters differ between Cornet and NMF and therefore could not be initialized identically.

### Applications to real tumor data

The following tumor WGS datasets were analyzed: colorectal cancers from PCAWG and GEL, oral/oropharyngeal cancers from GEL, B-cell non-Hodgkin’s lymphomas from PCAWG, and bladder cancers from PCAWG and GEL combined. For GEL data, only fresh-frozen samples were used, and samples with inconsistent tumor type annotations across multiple metadata sources were removed. For each dataset, samples with mismatch-repair deficiencies (MMRD) and/or POLE-exo mutations were identified by inspecting the exposures of SBS signatures known to be associated with MMRD and/or POLE-exo mutations through a refitting analysis (as in [11]) and removed beforehand. Additional outliers were identified with the preprocessing module in MuSiCal and removed. Cornet-derived *de novo* signatures are provided in the **Supplementary Tables**.

## Data availability

Mutation count matrices of PCAWG samples were downloaded from https://www.synapse.org/Synapse:syn11726601/files/. Mutation count matrices of GEL samples were obtained from the Supplementary Tables of [15]. COSMIC signatures were downloaded from https://cancer.sanger.ac.uk/signatures/. Signal signatures were obtained from the Supplementary Tables of [15]. Data from the National Genomic Research Library (NGRL) used in this research are available within the secure Genomics England Research Environment. Access to NGRL data is restricted to adhere to consent requirements and protect participant privacy. Data used in this research include clinical information and variants of potential clinical significance in cancer samples (accessed from the tables cancer_analysis and cancer_tier_and_domain_variants in LabKey, respectively). Access to NGRL data is provided to approved researchers who are members of the Genomics England Research Network, subject to institutional access agreements and research project approval under participant-led governance. For more information on data access, visit: https://www.genomicsengland.co.uk/research.

## Code availability

Cornet is implemented in the Python package Sonata, available at https://github.com/parklab/Sonata. Cornet is also integrated within the Python package MuSiCal for mutational signature analysis, available at https://github.com/parklab/MuSiCal.

## Author contributions

HJ conceived the study, proposed correlated NMF as the key methodology, supervised and contributed to algorithm development, established how to apply the algorithm to cancer data and interpret its output, integrated the algorithm into the MuSiCal framework, performed all benchmarking and cancer data analyses presented in this work, and designed and compiled the figures. BG formulated the Cornet model by refining correlated NMF, derived the mathematical framework, developed and implemented the algorithm and software, and performed initial benchmarking and data analysis. HJ and BG wrote the manuscript. DG tested the software and commented on the manuscript. DCG helped frame the research problem, provided conceptual guidance, and assisted with data access. PJP supervised the study, provided conceptual guidance, and revised the manuscript. All authors read and approved the final manuscript.

## Acknowledgements

This work was funded by grants from the National Institutes of Health (R01CA269805, R01HG012573, and UM1DA058230 to PJP). The funders had no role in study design, data collection and analysis, decision to publish or preparation of the paper. We gratefully acknowledge the participants of the National Genomic Research Library (NGRL), whose contributions made this research possible. Secure access to the NGRL under project ID 1087 was provided by Genomics England, which delivers the NGRL in partnership with NHS England, and is wholly owned by the UK Department of Health and Social Care. The NGRL contains participants’ health data collected by the NHS as part of their care, along with samples and data from their participation in research, for which fully informed consent has been obtained. This includes genomic and clinical data provided through the NHS Genomic Medicine Service, as well as data obtained through research studies, including the 100,000 Genomes Project and the Generation Study, both of which are delivered in partnership with the NHS, and from other research cohorts involving external collaborators.

## Supplementary Information

## Supplementary Methods

### Derivation of the Cornet objective and optimization algorithm

Here we provide a detailed derivation of the Cornet algorithm. Our derivation follows a similar strategy as [1], but the underlying generative model and optimization objective differ in several aspects (see **Methods**).

Let *V* be the number of mutation types, *K* the number of signatures, *D* the number of samples, and *m* the embedding dimension of the signature and sample embeddings *L* ∈ ℝ^*m×K*^ and *U* ∈ ℝ^*m×D*^. Let 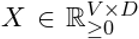 denote the mutation count matrix, 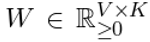 the signature matrix, and 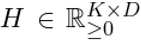 the exposure matrix. For signature offsets *α* ∈ ℝ^*K*^, sample offsets *β* ∈ ℝ^*D*^, variance parameter *σ*^2^ *>* 0, and the identity matrix *I*_*m*_ ∈ ℝ^*m×m*^, recall the generative model of Cornet:

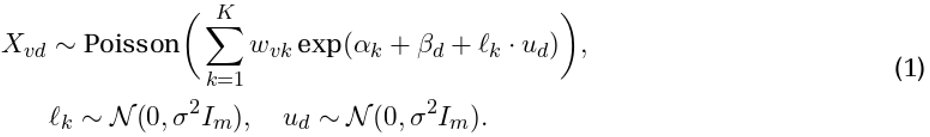

Thus, Cornet has the same Poisson observation model as KL-NMF,

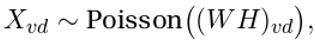

for which the negative log-likelihood is given, up to terms depending only on *X*, by the generalized Kullback–Leibler divergence

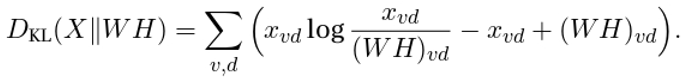

The key difference is that in KL-NMF the exposure matrix *H* is optimized directly, whereas in Cornet the exposures are parameterized as

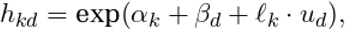

and Gaussian priors are placed on the latent embeddings in order to model the covariance structure in the exposures. Hereafter, we use NMF to refer specifically to KL-NMF with the standard multiplicative-update algorithm of [2].

### MAP derivation of the Cornet objective

We estimate the model parameters (*W, α, β, σ*^2^) and the latent embeddings (*L, U*) by maximizing the joint log-density

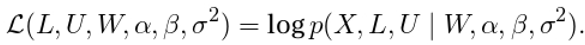

This corresponds to maximum a posteriori (MAP) estimation of the latent embeddings together with joint optimization of the remaining model parameters. As discussed in **Methods**, this formulation is closely related to the mean-field variational inference approach of [1] with independent point-mass variational distributions, but here we work directly with the MAP objective.

Using

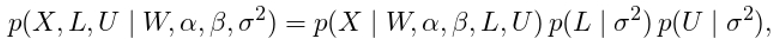

the objective function can be written as

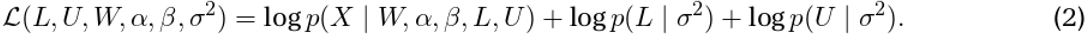

Assuming conditionally independent mutation counts and embeddings as in (1), we obtain

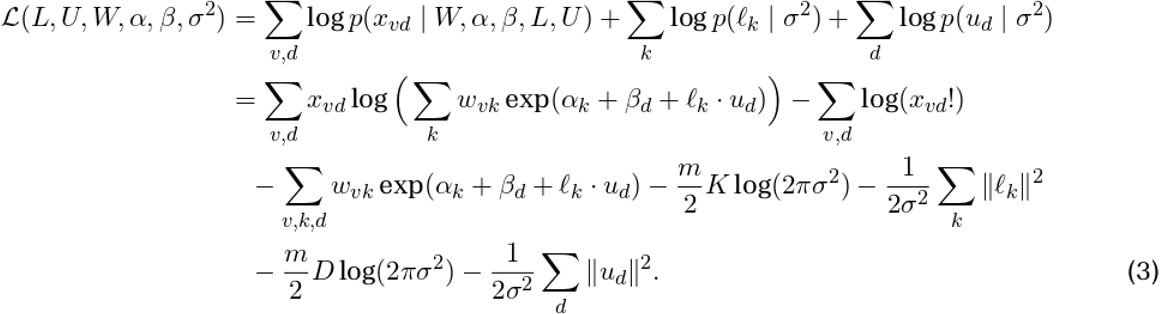

Discarding terms that depend only on *X*, this simplifies to

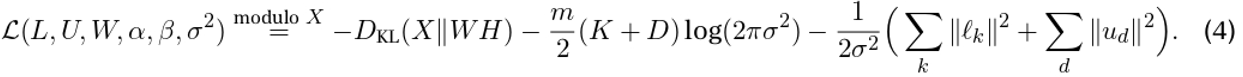

Thus, Cornet can be viewed as a version of NMF with an additional factorization of the exposure matrix to model correlations, together with a regularization term induced by the Gaussian priors on the latent embeddings.

### Coordinate ascent optimization

We optimize (4) by coordinate ascent. To handle the non-separable logarithmic term, we introduce a joint auxiliary function in the spirit of standard NMF auxiliary-function methods [1–5]. During optimization, we enforce the constraint that signatures are normalized, i.e.,

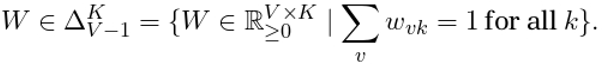

Let

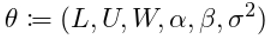

denote the full collection of optimization variables, and let *θ*^(*n*)^ be the current iterate. Briefly, the auxiliary-function principle proceeds by constructing a function ℒ_aux_(*θ, θ*^(*n*)^) such that

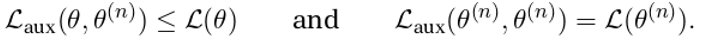

Maximizing this lower bound with respect to *θ* in each iteration guarantees that the objective value does not decrease:

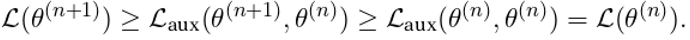

To construct such a bound, let

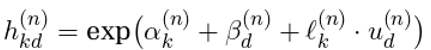

denote the current estimate of the exposure of signature *k* in sample *d*. For every *v, k*, and *d*, define

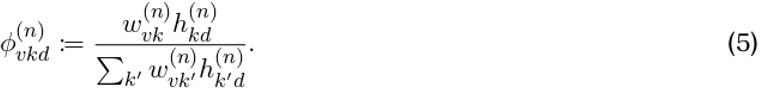

By construction, 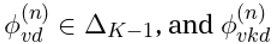 can be interpreted as the current estimate of the probability that a mutation of type *v* in sample *d* originated from signature *k*. Jensen’s inequality then implies

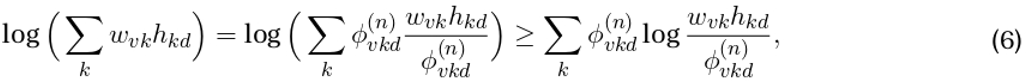

with equality at *θ* = *θ*^(*n*)^.

Substituting (6) into (4), we obtain the auxiliary function

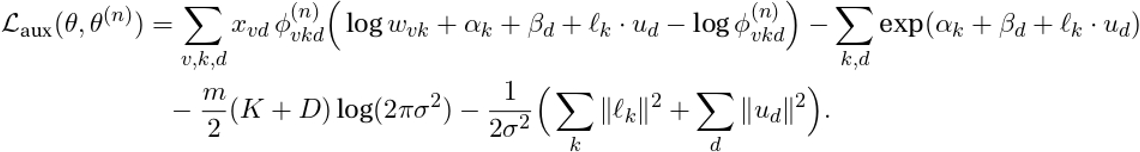

That is, ℒ_aux_(*θ, θ*^(*n*)^) is a lower bound on ℒ (*θ*), tight at *θ* = *θ*^(*n*)^. The Cornet algorithm updates *θ* by maximizing ℒ_aux_(*θ, θ*^(*n*)^) at each iteration.

#### Updating the latent embeddings

For fixed *θ*^(*n*)^, the terms of ℒ_aux_(*θ, θ*^(*n*)^) that depend on a single signature embedding *ℓ*_*k*_ are

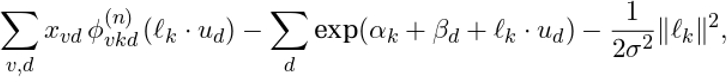

with gradient

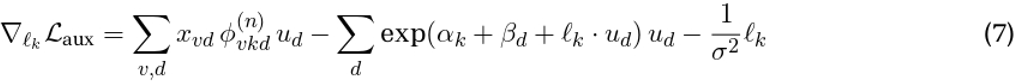

and Hessian

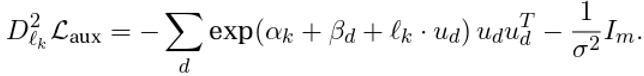

In particular, the Hessian is negative definite and maximizing ℒ_aux_ with respect to *ℓ*_*k*_ is a concave optimization problem.

Analogously, the terms of ℒ_aux_(*θ, θ*^(*n*)^) that depend on a single sample embedding *u*_*d*_ are

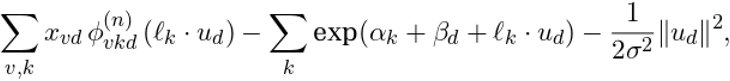

with gradient

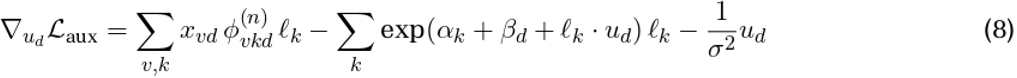

and Hessian

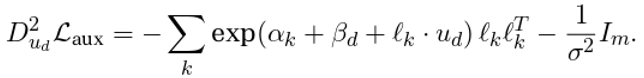

Thus ℒ_aux_ is also concave in each *u*_*d*_. In practice, we maximize ℒ_aux_ with respect to the embeddings using the `Newton-CG` algorithm implemented in `scipy.optimize.minimize` [6], although any suitable optimization method could be used.

#### Updating the model parameters

Maximizing ℒ_aux_(*θ, θ*^(*n*)^) with respect to each signature *w*_*k*_ ∈ Δ_*V* −1_ yields a closed-form update. Using the method of Lagrange multipliers, there exists *λ*_*k*_ ∈ ℝ such that

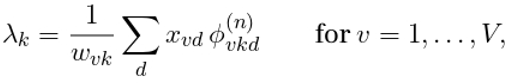

that is,

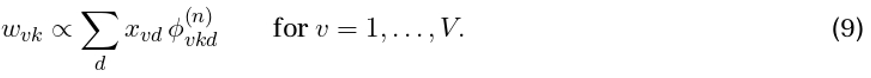

Because 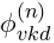 captures the relative contribution of signature *k* to mutation type *v* in sample *d*, (9) aggregates all mutations of type *v* currently attributed to signature *k*, after which normalization across mutation types defines the updated signature.

There are also closed-form maximizers of ℒ_aux_(*θ, θ*^(*n*)^) with respect to the signature offsets *α*_*k*_, the sample offsets *β*_*d*_, and the variance parameter *σ*^2^. First,

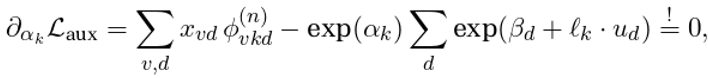

which yields

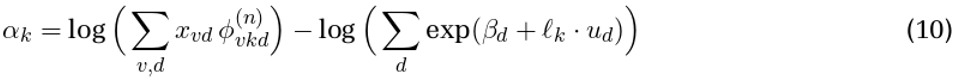

for all *k* = 1, …, *K*. Second,

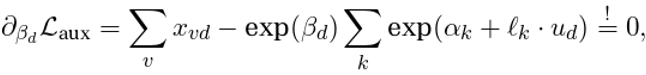

which gives the update rule

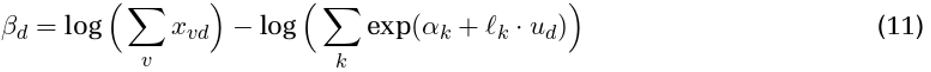

for all *d* = 1, …, *D*. Finally,

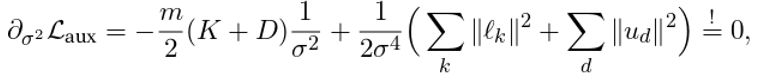

which yields

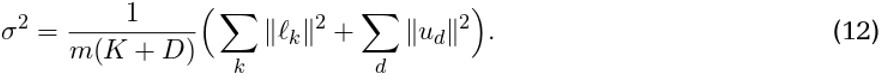

In [1], the formulas corresponding to (7), (8), and (11) are (11.30), (11.31), and (11.32), respectively. That work does not include signature offsets because the signatures are not normalized. In addition, [1] uses a fixed prior variance of 1 for the sample embeddings and treats the variance *σ*^2^ of the signature embeddings as a hyperparameter.

### Summary of the Cornet algorithm

The Cornet algorithm derived above is summarized in Algorithm 1. ConvexOptimizer denotes any unconstrained convex optimization algorithm. In practice, we do not compute and store the entire tensor *φ* at each iteration, but only the reconstruction *W H*, since all expressions involving *φ* that appear in the update rules can be rewritten using either

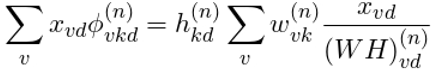

Or

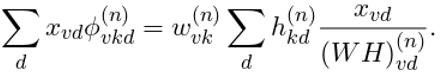

#### Algorithm 1

Cornet

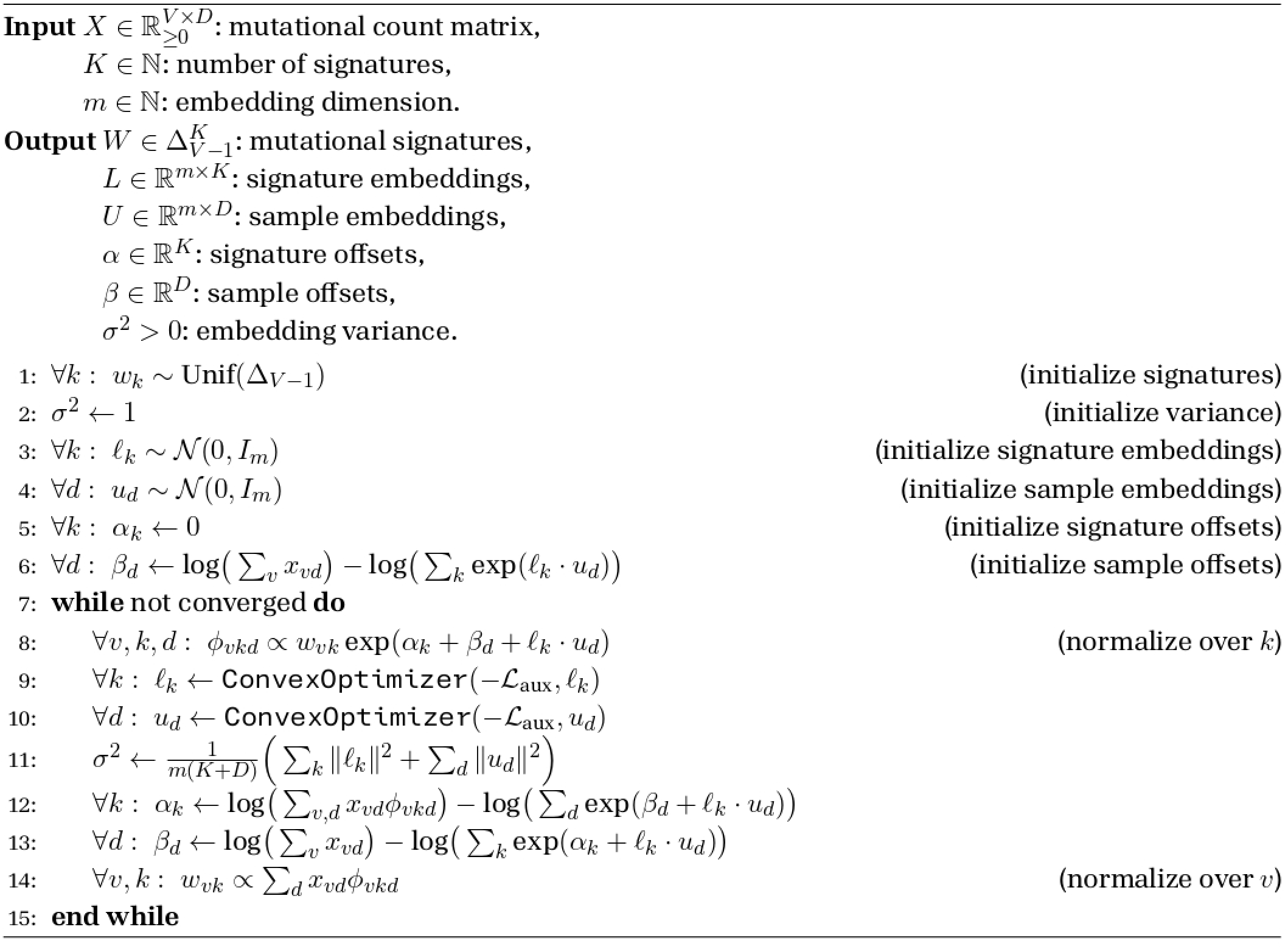

### Regularization and input scaling

In this section, we show that the scale of the mutational count matrix is closely related to the effective regularization strength in Cornet and therefore influences the sparsity of the fitted exposures. More specifically, after eliminating the sample offsets, Cornet with an arbitrary input matrix can be interpreted as a weighted version of Cornet on normalized count profiles.

The objective function (4) shows that Cornet can be viewed as a version of NMF with an additional low-dimensional factorization of the exposure matrix to model correlations, together with a regularization term induced by the Gaussian priors on the latent embeddings. Let

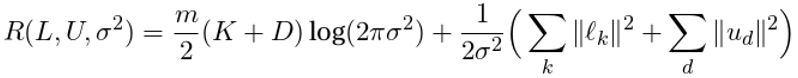

denote the regularization term. Then

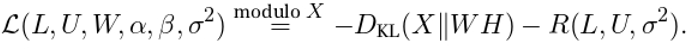

To make the effect of the scale of the mutational count matrix more explicit, we eliminate the sample offsets *β*_*d*_ and rewrite the objective in terms of normalized count profiles. Let

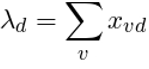

be the total mutational burden of sample *d*. Plugging the optimal sample offsets from (11) into the exposures *h*_*kd*_ yields

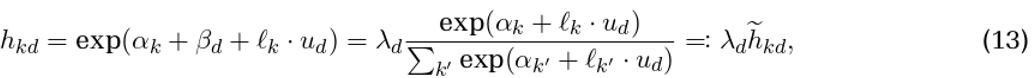

where 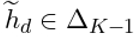. Thus, the sample offsets induce a softmax normalization of the log-exposures, followed by rescaling by the sample-specific mutation burden.

Let 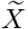 denote the normalized count matrix with 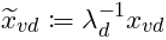, and define the weighted Kullback– Leibler divergence

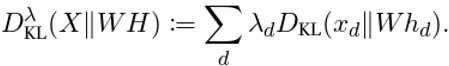

After eliminating the sample offsets via (13), the objective can be rewritten as

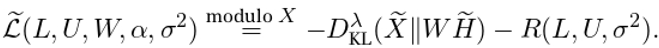

This rewriting shows that Cornet with an arbitrary input matrix *X* is equivalent to a weighted version of Cornet on the normalized count matrix 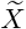, where the sample-specific weights are given by the total mutational burdens *λ*_*d*_. In particular, the quantities *λ*_*d*_ determine how strongly the reconstruction term contributes to the objective function relative to the regularization term.

This perspective provides a heuristic explanation for the effect of input scaling on the sparsity of the fitted exposures. Empirically, we observed that when the reconstruction term is weighted more strongly, i.e., for larger mutational burdens *λ*_*d*_, the norms of the embeddings can grow larger, which leads to sparser exposures. Conversely, when the regularization term is weighted more strongly, i.e., for smaller *λ*_*d*_, the embeddings are pushed more strongly toward the origin, leading to flatter exposures. Heuristically, the effect of the embedding norms on the softmax-normalized exposures is analogous to that of a temperature parameter. Through the coupled factorization problem, these changes in exposure structure can in turn affect the fitted signatures, but this relationship is more indirect. Empirically, we observed that sparser exposures were often accompanied by flatter signatures, and vice versa.

A related insight is obtained by plugging the optimal update

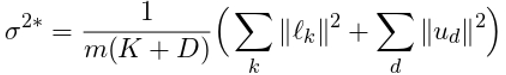

back into the regularization term. This yields

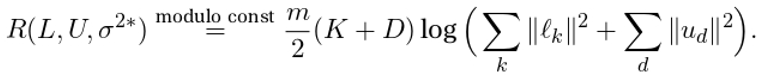

Thus, learning *σ*^2^replaces a fixed quadratic penalty by an adaptive logarithmic penalty on the total embedding norm. This is consistent with the discussion above: when the reconstruction term is weighted more strongly, larger embedding norms are favored, which in turn increases the learned variance *σ*^2^ and weakens the effective shrinkage toward the origin. Conversely, when the reconstruction term is weighted less strongly, the embeddings are pushed more aggressively toward zero, leading to smaller learned variance and flatter exposures.

Throughout the paper, we scaled the input SBS count matrices such that the average tumor mutational burden across samples was between 10*V* and 40*V*, where *V* is the number of mutation types (i.e., *V* = 96 for SBS analysis).

## Supplementary Figures

**Supplementary Figure S1.**
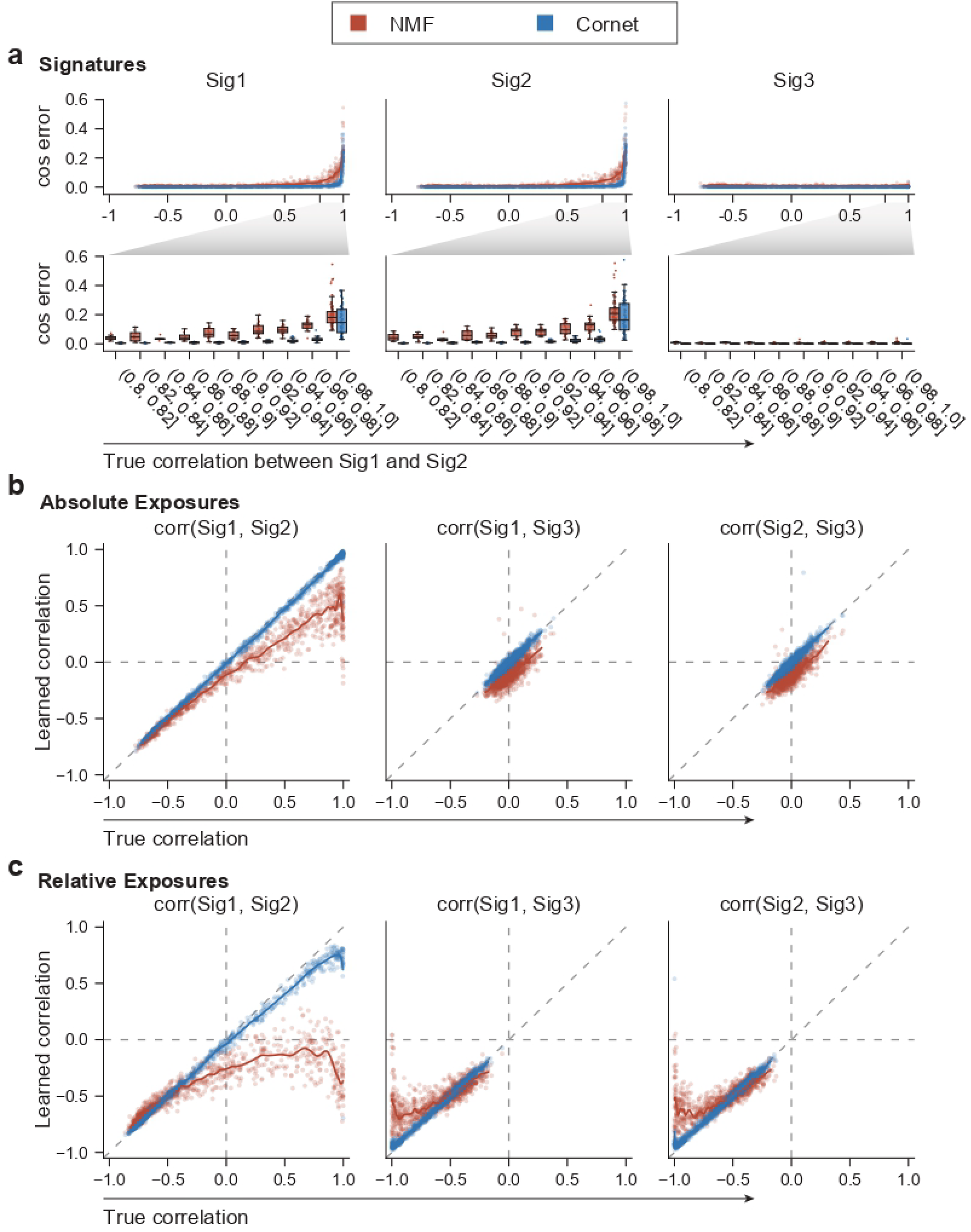
Benchmarking the performance of Cornet against NMF in simulated datasets. Synthetic mutation count matrices were generated from three random mutational signatures Sig1, Sig2, and Sig3. Correlations at varying strengths were introduced between Sig1 and Sig2, while Sig3 was independent of Sig1 and Sig2. **(a)** Accuracy of signature discovery, comparing Cornet with NMF solutions. Cosine errors between recovered and ground truth signatures are shown at different correlation levels. Results at strong correlations (*>*0.8) were enlarged and shown in the bottom row. **(b)** Accuracy of correlation structure inference, comparing Cornet with NMF solutions. Inferred correlations were plotted against the ground truth correlations for each pair of the three signatures. Pearson correlation coefficients using absolute exposures are shown. **(c)** Same as (b) but using L1-normalized relative exposures. Note that this normalization procedure induces strong negative correlations between Sig1/2 and Sig3 in the normalized ground-truth exposures.

**Supplementary Figure S2.**
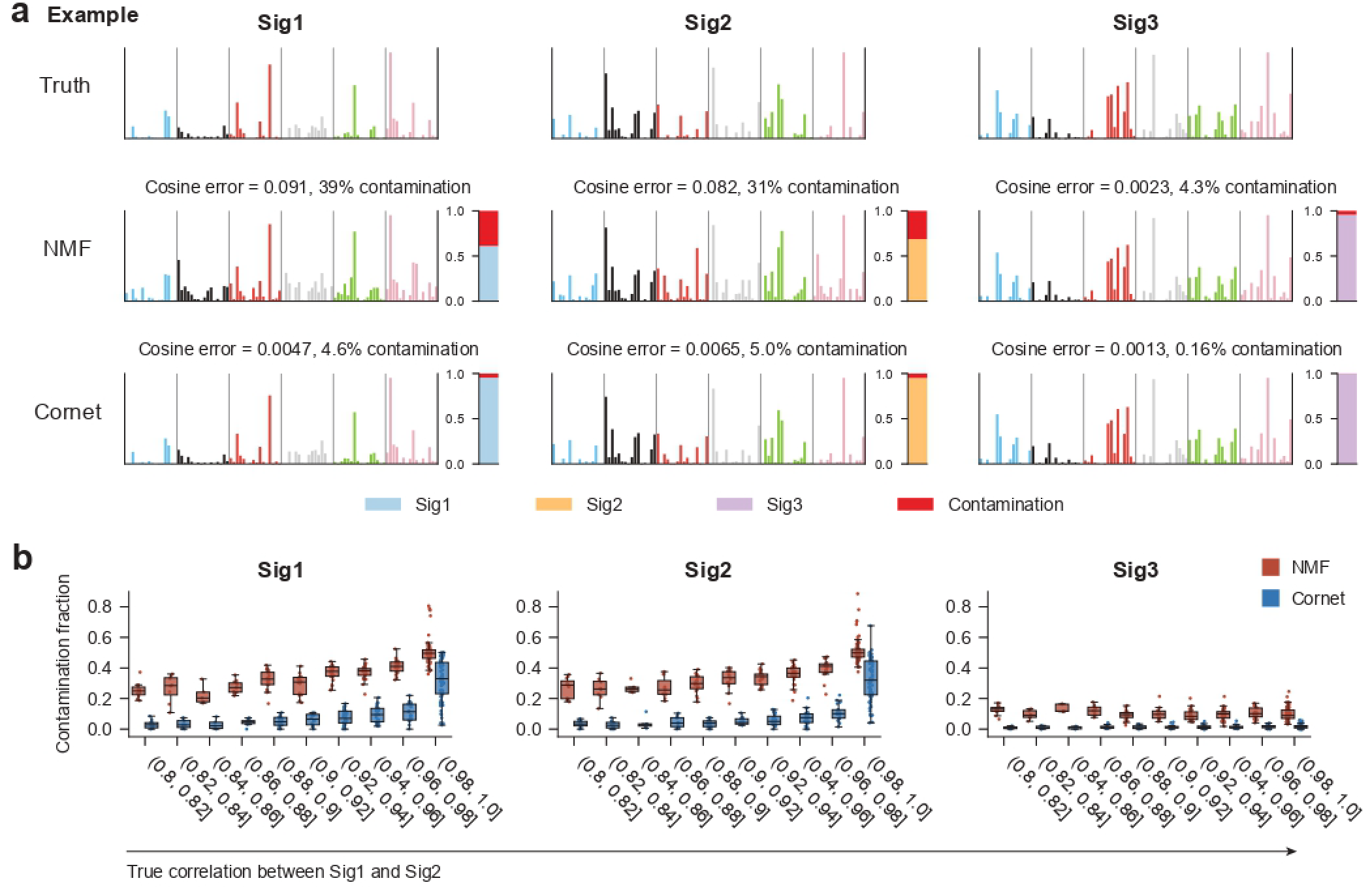
Cornet-derived signatures are less contaminated in simulated benchmarks even under strong correlations. **(a)** Simulated example demonstrating that seemingly small cosine errors can correspond to large contaminations. Top: ground truth signatures. Middle: NMF solutions together with bar plots of NNLS weights obtained by decomposing the NMF solutions into ground truth signatures. Bottom: Cornet solutions together with bar plots of NNLS weights obtained by decomposing the Cornet solutions into ground truth signatures. **(b)** Contaminating weights at different correlation levels, comparing NMF and Cornet solutions.

**Supplementary Figure S3.**
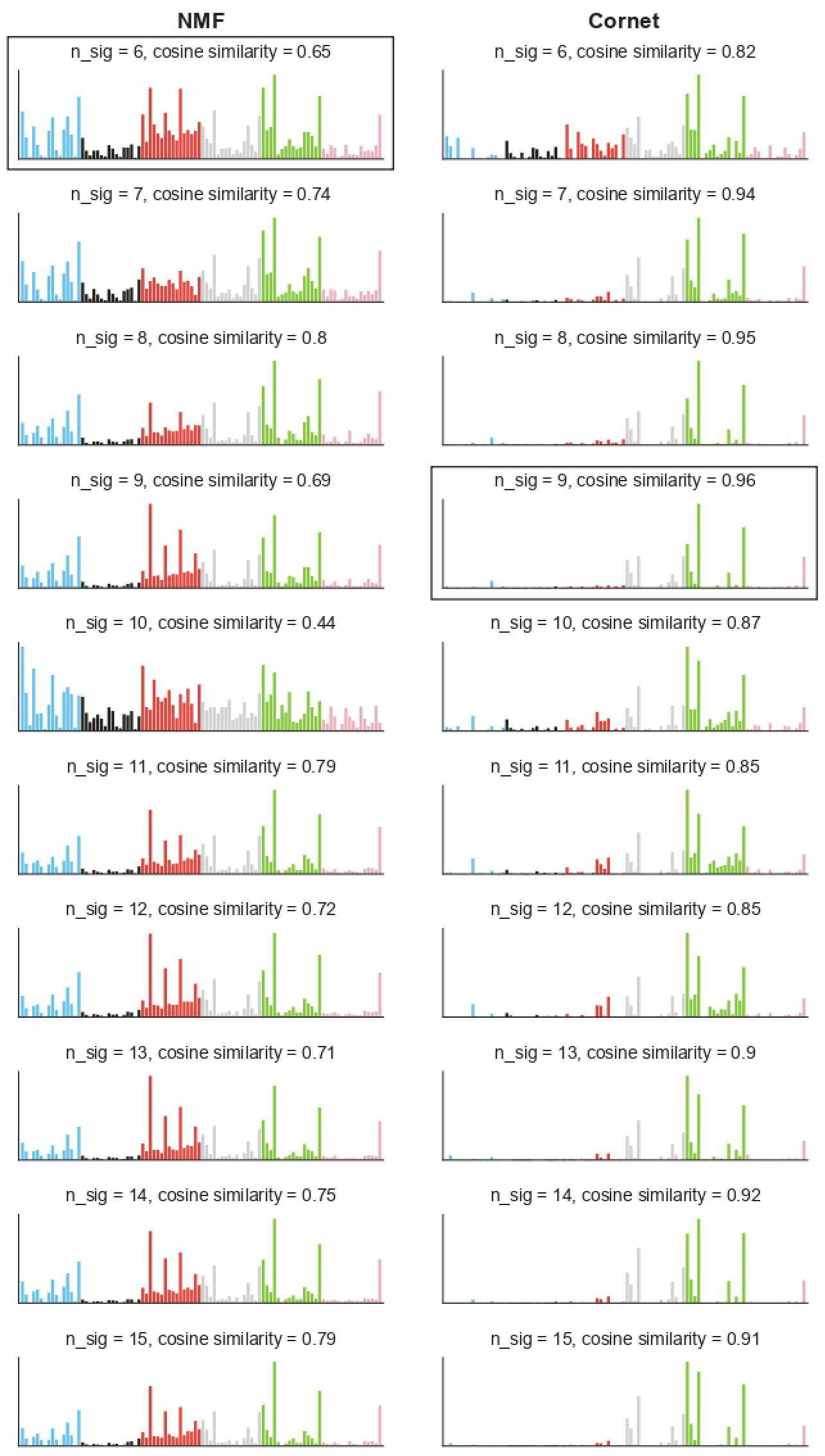
NMF failed to accurately discover the colibactin signature SBS88 from the small PCAWG colorectal cancer dataset in **Fig. 2** even when forced to return more signatures. Left: Top SBS88-matching NMF-derived *de novo* signatures for solutions at different numbers of signatures. Right: Top SBS88-matching Cornet-derived *de novo* signatures for solutions at different numbers of signatures. The signatures shown in **Fig. 2c** and **Fig. 2f** are highlighted with boxes.

**Supplementary Figure S4.**
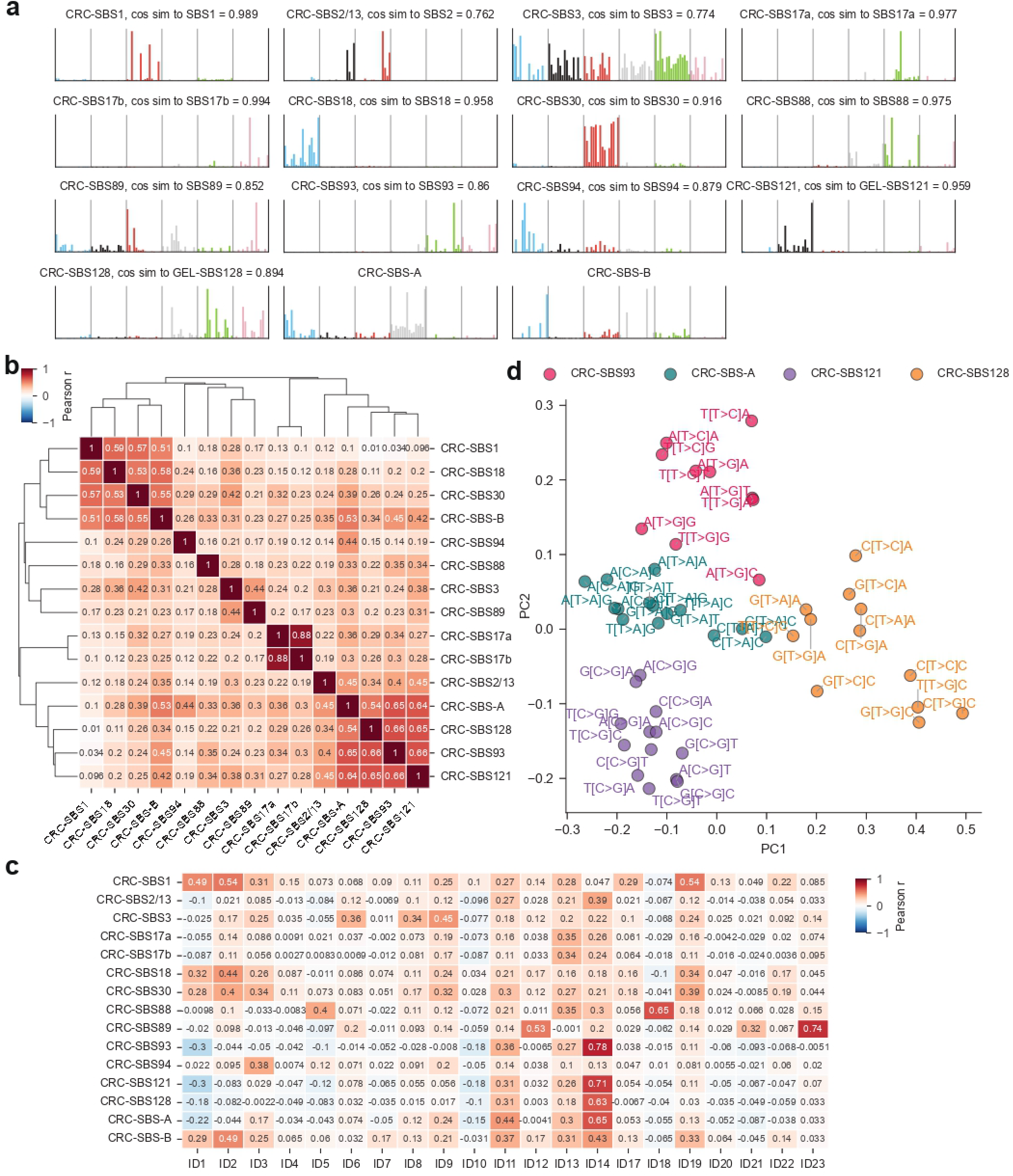
Additional information on the CRC analysis in Fig. 4. **(a)** All Cornet-derived *de novo* signatures discovered in CRC. **(b)** Pearson correlation coefficients between the exposures of *de novo* signatures. Signatures were clustered using hierarchical clustering. **(c)** Pearson correlation coefficients between the exposures of Cornet-derived *de novo* signatures and COSMIC indel signatures. **(d)** Validation of the split of COSMIC SBS93 using PCA. Top 50 samples with the largest total exposures of CRC-SBS93, CRC-SBS-A, CRC-SBS121, and CRC-SBS128 were analyzed. PCA with a cosine distance kernel was applied to the mutation count matrix, and the mutation types were plotted in the PC1-PC2 space. Further, each mutation type was assigned to a *de novo* signature based on which signature was most likely to have generated a mutation of that type. 49 of the 96 types were assigned to CRC-SBS93, CRC-SBS-A, CRC-SBS121, or CRC-SBS128, and included in the analysis.

**Supplementary Figure S5.**
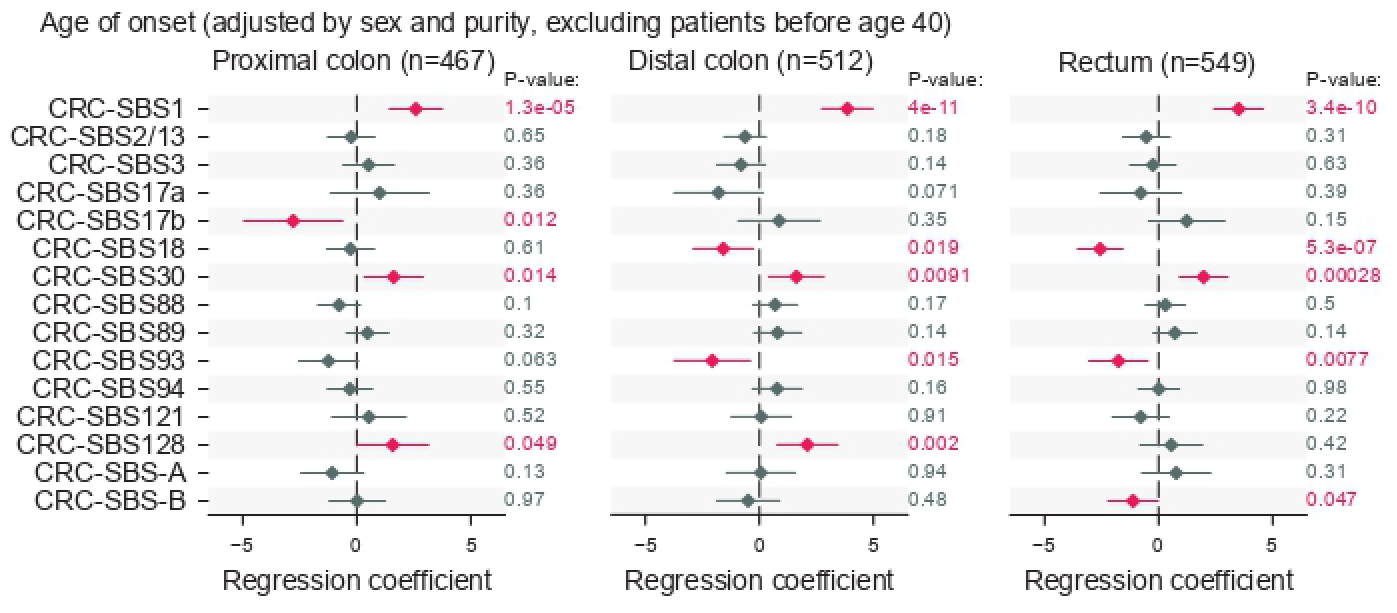
Same as **Fig. 4f**, except that patients younger than 40 years were excluded from the analysis.

**Supplementary Figure S6.**
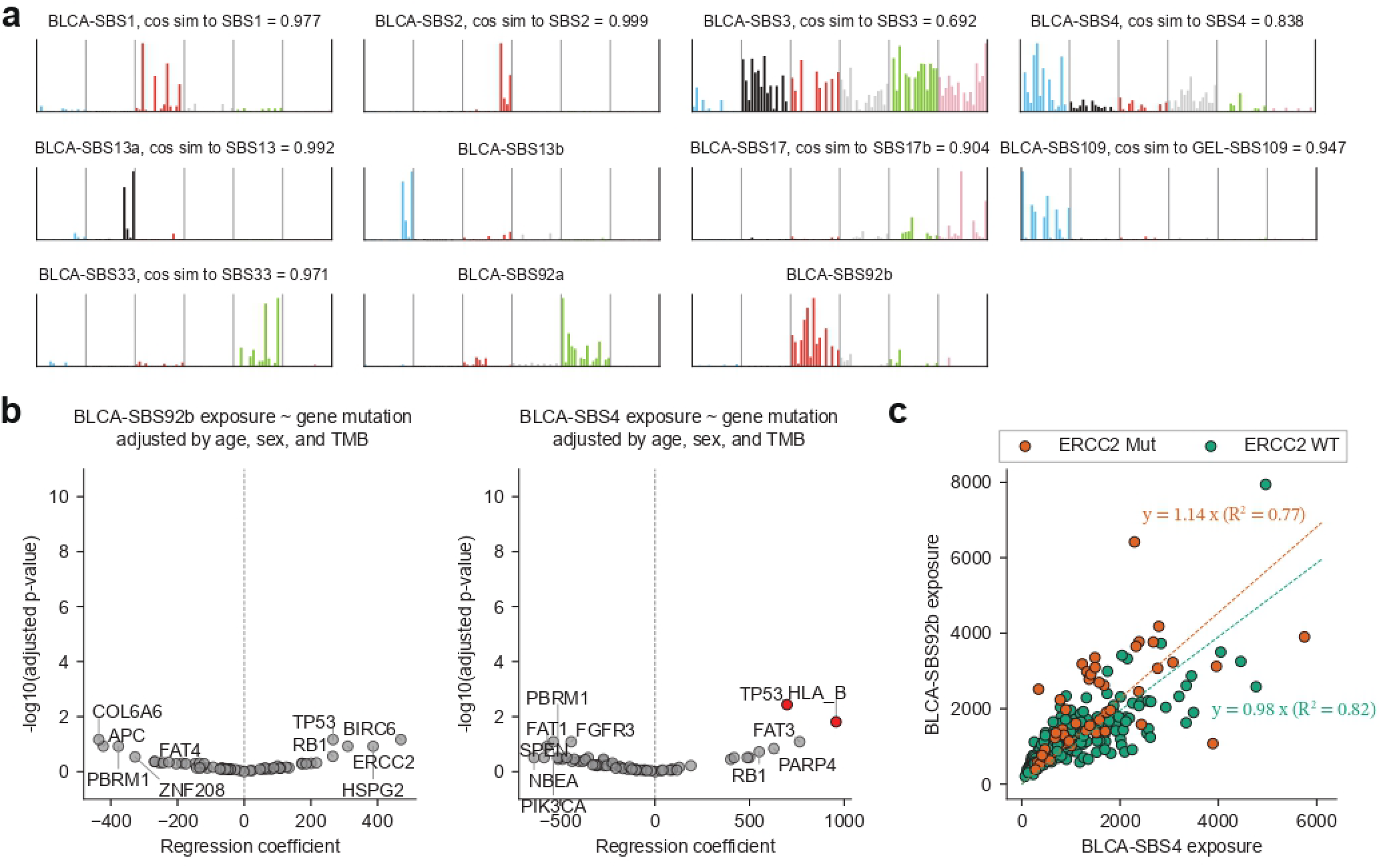
Additional information on the BLCA analysis in Fig. 5. **(a)** All Cornet-derived *de novo* signatures discovered in BLCA. **(b)** Volcano plots for the linear regression of BLCA-SBS92b (left) and BLCA-SBS4 (right) exposures against gene-level mutation status. Patient age, sex, and tumor mutational burden (TMB) were included as covariates in the regression. Genes with an adjusted p-value *<*0.05 are colored red. **(c)** Absolute exposures of BLCA-SBS92b vs. BLCA-SBS4 for ERCC2-widetype (WT) and ERCC2-mutant (Mut) samples. Linear regression slopes are annotated. One outlier sample, with a BLCA-SBS4 exposure more than four times higher than that of any other sample, was removed.

